# Reduced GABA transmission onto ventral tegmental area dopamine neurons underlies vulnerability for hyperactivity in a mouse model of Anorexia Nervosa

**DOI:** 10.1101/2024.03.14.585038

**Authors:** Fabien Ducrocq, Eileen Brouwer, Karlijn L. Kooij, Inge G. Wolterink-Donselaar, Lisa Drost, Jaimie Hak, Melissa Veendijk, Mieneke C. M. Luijendijk, Frank J. Meye, Roger A. H. Adan

## Abstract

Anorexia nervosa (AN) has the highest mortality among psychiatric diseases. Hyperactivity is a persistent symptom, which is difficult to control for patients and a major barrier to recovery as it interferes with weight gain. Alteration of mesolimbic dopamine transmission has been hypothesized as a critical factor for the development and maintenance of the disease and for hyperactivity. At what level the changes in dopamine occur in anorexic states and whether local mesolimbic neurocircuit plasticity is causally involved remains unclear. Especially the role of local GABA control over dopamine neurons, a powerful regulator of the dopamine system, in an AN context is unknown. We hypothesize that combining caloric restriction with exercise, such as in the activity-based anorexia (ABA) model, alters dopamine transmission via GABA disinhibition that, in turn, facilitates the expression of maladaptive behaviors such as hyperactivity.

Therefore, we characterized the impact of the ABA model on plasticity of the dopamine reward system. In ex-vivo brain slices of mice exposed to this model, ventral tegmental area dopamine (VTA_DA_) neurons displayed a higher firing frequency compared to control mice supporting that the midbrain dopamine system undergoes plasticity. This coincided with reduced GABAergic transmission on VTA_DA_ neurons. This reduction was at least in part attributable to local VTA GABA (VTA_GABA_) neurons. Indeed, VTA_GABA_ neurons were less excitable, displayed a lower firing frequency and a lower probability of release onto VTA_DA_ neurons. Restoring the excitability of VTA_GABA_ neurons via chemogenetic activation rescued mice from starvation, by decreasing running wheel activity.

In summary, we found that the anorexic state leads to dysregulation of VTA_GABA_ transmission on VTA_DA_ neurons that reinforces maladaptive behaviors such as excessive exercise. We uncovered a new mechanism linked to the disturbed dopamine system in ABA-exposed animals, identifying a hitherto unknown role of decreased local GABAergic control over VTA dopamine neuron output.

## INTRODUCTION

Anorexia nervosa (AN) is a metabo-psychiatric eating disorder characterized by a fear of gaining weight and self-starvation, leading to life-threatening weight loss [1, 2]. Although not part of the formal diagnostic criteria, increased drive for activity is an important aspect of this disorder, with up to 80% of AN patients engaging in excessive exercise and displaying restlessness [3, 4]. Hyperactivity interferes with weight gain and has been found to predict relapse and chronicity of AN and therefore represents a major barrier to recovery [5, 6].

The dopamine (DA) system has been implicated in the neurobiology underlying AN since the 70s [7] and has been strengthened by recent clinical evidence of dysfunction of the reward system in AN [8–12]. More precisely, recovered AN patients exhibit reduced levels of a DA metabolite in cerebral spinal fluid [13] and increased DA D2/D3 receptor binding in the ventral striatum [10], although the latter was not confirmed in another study [14]. Furthermore, AN patients display altered reward processing associated with differential responses in the ventral striatum depending on the rewarding stimulus [15], reduced VTA / susbtancia nigra to nucleus accumbens functional connectivity [16], increased responsiveness of reward circuits to food stimuli [11]. An altered effective connectivity was found in AN patients between the feeding system and the DA reward system when anticipating a sweet reward (i.e. in participants with AN, the ventral striatum directed effective connectivity to the hypothalamus whereas hypothalamus to the ventral striatum dynamics were observed in controls, [12]). These findings indicate that there are alterations in DA signaling in AN, but the precise nature and mechanism underlying these alterations remain unknown.

Animal studies support the link between the DA reward system and the regulation of energy homeostasis. DA transmission in the ventral tegmental area (VTA) is involved in energy homeostasis by regulating both exercise and feeding behaviors [17–22]. The activity-based anorexia (ABA) model is typically used to better understand the neurobiology of AN and evaluate potential treatment targets (for review see [23, 24]). Studies using the ABA model have shown that manipulation of dopamine transmission facilitates the development of behaviors that resemble AN symptoms, such as excessive exercise and weight loss, supporting the role of altered DA neuron activity in hyperactivity (for review see [25]). Mice without the dopamine transporter gene (DAT knockouts) are hyperdopaminergic and more susceptible to lose weight in the ABA model due to accelerated development of excessive running [26]. Chlorpromazine, a dopamine receptor 2 (D2) antagonist, was shown to reduce locomotor activity and prevent “self-starvation” in rats [27], a finding that was replicated with the D2/3 antagonists eticlopride and amisulpride [28]. Overexpression of D2 receptors in ventral striatum increased hyperactivity and the susceptibility to lose weight in the ABA model [29]. These rodent data align with patients with AN displaying hyperactivity in particular during the acute phase of the disease [3]. Strikingly, it is still unclear how environmental factors that drive negative energy balance, such as food restriction and the availability to exercise, impact on DA transmission.

Although alterations have been observed in the DA system of AN patients and manipulations of dopamine transmission have been shown to impact on AN-like symptoms in animals exposed to the ABA model, there remains a gap in understanding the specific alterations occurring in the dopamine system. Alterations in intrinsic properties of DA neurons may be causal to changed DA transmission, yet altered GABA transmission has been proposed as a key player in ABA vulnerability and maladaptive behaviors. Individuals with the greatest expansion of the GABAergic synapses on hippocampal and prefrontal pyramidal neurons were the most resilient to ABA exposure [30]. Interestingly, in a study using deep brain stimulation in the VTA, suppression of hyperactivity was attributed to increased VTA GABA activity in a mouse model of schizophrenia [31]. GABAergic inhibition of the dopamine system emerges as a candidate mechanism, given its potent regulatory influence on dopamine neuron output [32, 33]. However, to date, the impact of ABA-exposure on VTA_DA_ neurons and the involvement of GABA transmission in DA alterations is unknown. This study aimed at unravelling how exposure to the ABA model impacts on DA neurons in female mice and at identifying neuronal mechanisms involved. We hypothesize that hyperactivity of the midbrain DA system is involved in hyperactivity and body weight loss through local GABA disinhibition of VTA_DA_ neurons. To investigate this, we assessed 1) the consequences of the ABA model on DA neurons in the VTA, 2) the GABAergic origin of the VTA_DA_ neuronal alterations and 3) the influence of modulating GABA neurons on the vulnerability to the ABA model. This work contributes to a better understanding of the interactions between energy balance regulation, the newly-described VTA_GABA_ alterations and its impact on VTA_DA_ neurons. Ultimately, it leads to a deeper comprehension of AN etiology and suggests that targeting the VTA_GABA_ system has therapeutic potential.

## METHOD DETAILS

### Experimental model and subject details

In all experiments, naïve young adult female mice were used (18–30g, 2-5 months). Before experiments started, all mice were group-housed in eurostandard type II cages (n=2–6, Tecniplast) in a 12:12 light/dark cycle (lights on at 07:00 a.m.) at 22 ± 2 °C (60–65% humidity) and properly handled. Unless otherwise specified, animals had access to ad libitum water and lab chow (3.61 kcal/g, CRM(E), 801730, Special Diets Services) and cage enrichment. The following mouse lines were used in this study: C57Bl6J (Jax#664), Pitx3-GFP (Jax#041479) and VGAT-Cre (Jax#028862) mice. Animals were bred in house but originated from the Jackson Laboratory. Homozygous VGAT-cre females were crossed to homozygous or heterozygous Pitx3-GFP males in order to generate double transgenic animals. Experiments were approved by the Animal Ethics Committee of Utrecht University and the Dutch Central Authority for Scientific Procedures on Animals (CCD#AVD1150020198686), and were conducted in agreement with the Dutch law (Wet op de Dierproeven, 2014) and the European regulations (Guideline 86/609/EEC). Every effort was made to minimize suffering and reduce the number of animals used.

### Activity-Based Anorexia (ABA) model and *ad libitum* running/feeding setup

Mice were single housed and randomly attributed, based on body weight (0.01gr resolution), to one of three groups: wheel running (WR), activity-based anorexia (ABA) and food restricted (FR). To reach stable running wheel activity (RWA), all mice were habituated for 10 days with ad libitum access to food and a running wheel in the unlocked (WR and ABA groups) or locked (FR) configuration. At ABA0, food was removed from FR and ABA mice 3 h after the start of the dark cycle. On all following days, food was provided for the first 3h of the dark cycle. For mice in the WR group, food was measured at dark onset daily, but not removed. Body weight was measured in all mice immediately prior to the start of the dark cycle (before food presentation). RWA was continuously collected and analyzed using a Cage Registration Program (Department Biomedical Engineering, UMC Utrecht, The Netherlands). Food anticipatory activity (FAA) was defined as RWA recorded during the last 6h of the light phase (unless otherwise indicated). We defined as humane endpoint (HEP) more than 20% loss of initial body weight that were not recovered following food exposure. The 20% loss of initial body weight was used as cut off for survival analyses. Mice were habituated to intraperitoneal (i.p.) injections (100ul of saline/10g of mice) for at least three days before the onset of the behavioral paradigm (ABA 0). In Fig.4, the habituation phase described above was followed by an *ad libitum* protocol (for both food and running) in order to assess the dose-dependent effect of CNO on feeding, RWA and variation of body weight.

### CNO administration

For chemogenetic experiments, Clozapine N-oxide (CNO) dihydrochloride (HB6149, Hello Bio) was dissolved in saline and injected i.p. at different doses (from 0.04mg/kg to 2mg/kg) with a volume of 100ul/10gr of mice. For chemogenetic inhibition, CNO was injected once a day, 5 hours before dark onset, at 0.5mg/kg. For chemogenetic activation, CNO was injected twice a day in order to have a better coverage of the period in which CNO was in the system (5 hours before dark onset and after feeding) at 0.25mg/kg. The measurements of body weight and food intake were performed at dark onset and after feeding. FAA was defined as the 5 hours following CNO injections. For CNO dose determination in the *ad libitum* feeding / running protocol, different CNO doses were randomized and injected at dark onset daily (ranging from 0.25 to 2mg/kg for inhibition and from 0.04 to 0.625mg/kg for activation). The measurements of body weight and food intake were performed at injection time and 3 hours later. RWA activity was measured as the sum of revolutions performed during these 3 hours.

### Stereotaxic Surgeries for Viral Injections

Stereotaxic injections of viral vectors were performed with a stereotaxic apparatus (UNOB.V. model 68U025) under anesthesia with ketamine (75mg/kg i.p.; Narketan, Vetoquinol), dexmedetomidine (1mg/kg i.p.; dexdomitor, Vetoquinol) and carprofen (5mg/kg subcutaneously or s.c., Carporal) injected 30 minutes before surgery. Lidocaine (0.1ml; 10% in saline; B. Braun) was injected s.c. on the skull before surgery. Animals were kept on a heat pad during surgery (≈37 °C). Viral injections were done using a 31G metal needle (Coopers Needleworks) attached to a 10μl Hamilton syringe (model 801RN) via flexible tubing (PE10, 0.28mm ID, 0.61mm OD, Portex) and an automated pump (UNO B.V. model 220). Mice were injected bilaterally with a volume of 300nl per side at an injection rate of 100nl/min with pAAV5-hSyn-DIO-hM4D(Gi)-mCherry (44362-AAV5, AddGene), pAAV-hSyn-DIO-hM3D(Gq)-mCherry (44361-AAV5, AddGene), pAAV-EF1a-double floxed-hChR2(H134R)-mCherry-WPRE-HGHpA (20297-AAV5, AddGene) or pAAV-hSyn-DIO-mCherry (50459-AAV5, AddGene) at titer 3.3*10^12gc/ml in the VTA (AP:-3.2, ML: 1.6, DV: -4.8 from bregma, angle 15°, Paxinos and Franklin). The injection needle was retracted 100μm, 9min after the infusion and withdrawn completely 10min after the infusion. The incision was sutured (V926H, 6/0, VICRYL, Ethicon) and animals were subcutaneously injected with the antagonist atipamezole (50mg/kg; Atipam, Dechra) and 1ml of saline and left to recover on a warm plate of 36°C. Carprofen (0.025mg/L) was provided in the drinking water all along post-surgery welfare monitoring. Experiments were performed 3-6 weeks after AAV stereotaxic injection. Injection sites were checked in all animals by preparing sagittal sections of 250µm for electrophysiology or 50µm after chemogenetic *in vivo* experiments. As in all mice the VTA was hit, but often with some variable spread to surrounding GABA neurons, we refer to these neurons as midbrain GABA neurons.

### Slice Preparation, Cell-attached and Whole-Cell Patch Clamp Recording

Mice were anesthetized with isoflurane at 12 p.m. (corresponding to dark onset) and then rapidly decapitated. The brain was quickly removed and coronal slices (250µm) were prepared as described previously [34]. Briefly, an ice-cold carbogenated cutting solution was used to prepare slices which then were transferred in the same solution for 5 min at 37°C. Next, they were stored at room temperature in an incubation medium for at least 60min. During recordings, slices were transferred to a recording chamber and perfused continuously at 1.5-2 mL/min with oxygenated artificial cerebrospinal fluid (ACSF). All solutions were prepared with pH≈7.35 and ≈300-310mOsm. Cells were visualized with an epifluorescent microscope (Examiner A1, Zeiss). Patch pipettes (2.7–4 MΩ for whole-cell patch-clamp and 6.0 and 6.5 MΩ for cell attached mode) were pulled from borosilicate glass capillaries (GC150-10, Harvard apparatus, UK) with a pipette puller (P-97, Sutter Instruments). Electrophysiological recordings were performed using a Axopatch 200B amplifier (Molecular Devices) and acquired using a Digidata 1322A digitizer (Molecular Devices), sampled at 20 kHz for current clamp, voltage-clamp recordings and cell-attached recordings, and low-pass filtered at 1 kHz. All data acquisitions were performed using pCLAMP 9.2 software (Molecular Devices). Light pulses were delivered with a pE-300 (CoolLed, UK illumination system). Only cells that maintained a stable access resistance (< 30MΩ, less than 20% increase) were included in analyses. In Pitx3-GFP mice, GFP-positive dopaminergic cells were recorded within the VTA. In VGAT-cre/Pitx3-GFP mice injected with cre-dependent AAV-ChR2-mCherry, GFP+/mCherry-dopaminergic cells were recorded within the VTA. In VGAT-cre/Pitx3-GFP mice injected with cre-dependent AAV-mCherry, mCherry+/GFP-GABAergic cells were recorded within the VTA.

For cell-attached recordings of dopaminergic neurons, we used filtered extracellular medium in the glass pipette and pipette resistance during recording was between 20-300MΩ. Recordings were made in voltage clamp gap free mode. Cells were clamped at 0mV and kept in this configuration for at least 10min. For recordings of basic intrinsic properties, we used the potassium-gluconate internal solution which consisted of (in mM) potassium gluconate 139; HEPES 10; EGTA 0.2; creatine phosphate 10; KCl 5; Na2ATP 4; Na3GTP 0.3; MgCl2 2, pH 7.35. To record basic intrinsic properties of VTA_DA_ and VTA_GABA_ neurons, cells were held in current-clamp mode in the absence of any current injections. A series of depolarizing current steps of 800ms starting at -105pA with 15pA increment were then applied. Cesium-chloride internal solution which consisted of (in mM) CsCl 139, NaCl 5, MgCl2 2, EGTA 0.2, HEPES 10, creatine phosphate 10, Na2-ATP 4, Na3-GTP 0.3, spermine 0.1, pH 7.35 was used to record GABA-mediated inhibitory transmission and paired-pulse ratios (PPR). IPSCs were recorded in the presence of 10µM CNQX (6-cyano-7-nitroquinoxaline-2,3-dione, Tocris 1045) and 50µM D-AP5 (D-2-amino-5-phosphonovalerate, Tocris 0106). Neurons were voltage clamped at -70mV. Recordings of spontaneous (s)IPSCs were performed at least 10min after patch for 1min. PPR recordings of VTA-driven GABA_A_R-mediated currents were performed in the presence of 10µM CNQX and 50µM D-AP5. GABA_A_R mediated currents were measured in response to optostimulation of VTA_GABA_ terminals using two pulses with 50, 100 and 200ms inter-pulse interval. The PPR was calculated by dividing the average amplitude of the evoked synaptic response (eIPSC) to the 2nd pulse by the amplitude of the eIPSC to the 1st pulse. Light pulses: 470 nm, 0.1 ms; 1-10mW. Data were analyzed offline using Clampfit 11.2 (MolecularDevices).

### Quantification and statistical analysis

Student’s t-tests (unpaired and paired) or equivalent non-parametric tests when data were not normality distributed (Mann-Whitney and Wilcoxon matched-pairs signed rank test) were used when two groups were compared. Welch’s correction was applied on unpaired t-test when variances were not equal. One-way ANOVA test or equivalent non-parametric tests when normality failed (Kruskal-Wallis test) were used when 3 groups were compared. Two-way ANOVA tests were used when 2 factors were present (Two-way repeated-measure ANOVA when one of the factors was a repeated-measure). Sidak’s post hoc test was applied when ANOVA showed a significant interaction or a significant main effect. Pearson’s Correlation Coefficient was used to measures the strength of the linear relationship between two variables. Log rank Mantel Cox test was used to compare survival curves. All statistical data were obtained using GraphPad Prism 7 (Graphpad Software). Statistical significance was *p < 0.05, **p < 0.01, ***p < 0.001 (#p<0.1 was used as an indication of a trend). All data are presented as means ±SEM. Details of the statistical analysis per figure are summarized in Tables S1 and S2 (main and supplementary figures respectively).

## RESULTS

### Mice exposed to the ABA model lose weight and develop excessive running wheel activity

We exposed female mice to the ABA model. The ABA protocol consisted of multiple days with a 3h feeding period at dark onset, combined with unlimited access to a running wheel (Fig.1A). Compared to FR (exposed to food restriction but with no wheel access) and WR (running wheel access and no food restriction) controls, ABA mice lost more body weight (Fig. 1B) while they ate a similar amount of food as FR mice and both groups ate less than WR controls (Fig. 1C). During the 5 days of ABA exposure, increased light phase RWA develops (Fig.1D). This maladaptive behavior, defined as food anticipatory activity (FAA), occurs the hours before food exposure at dark onset. Overall, ABA and wheel running (WR) animals display similar 24h running distances (Fig1.E). ABA mice decrease dark phase running activity, especially when food is available (Fig.1D), while FAA develops as running shifts to the light phase (Fig.1F). We found that body weight was positively associated with food intake (Fig.S1A) and negatively with FAA (Fig.S1B). Moreover, the animals displaying the highest RWA activity before ABA starts were modestly more vulnerable to develop FAA and to lose body weight (Fig.S1C-D), similarly to what has been previously found [35]. Altogether, these data show that in the ABA model the combination of reduced food intake and access to a running wheel leads to negative energy balance and produce AN-like symptoms.

**Figure 1.**
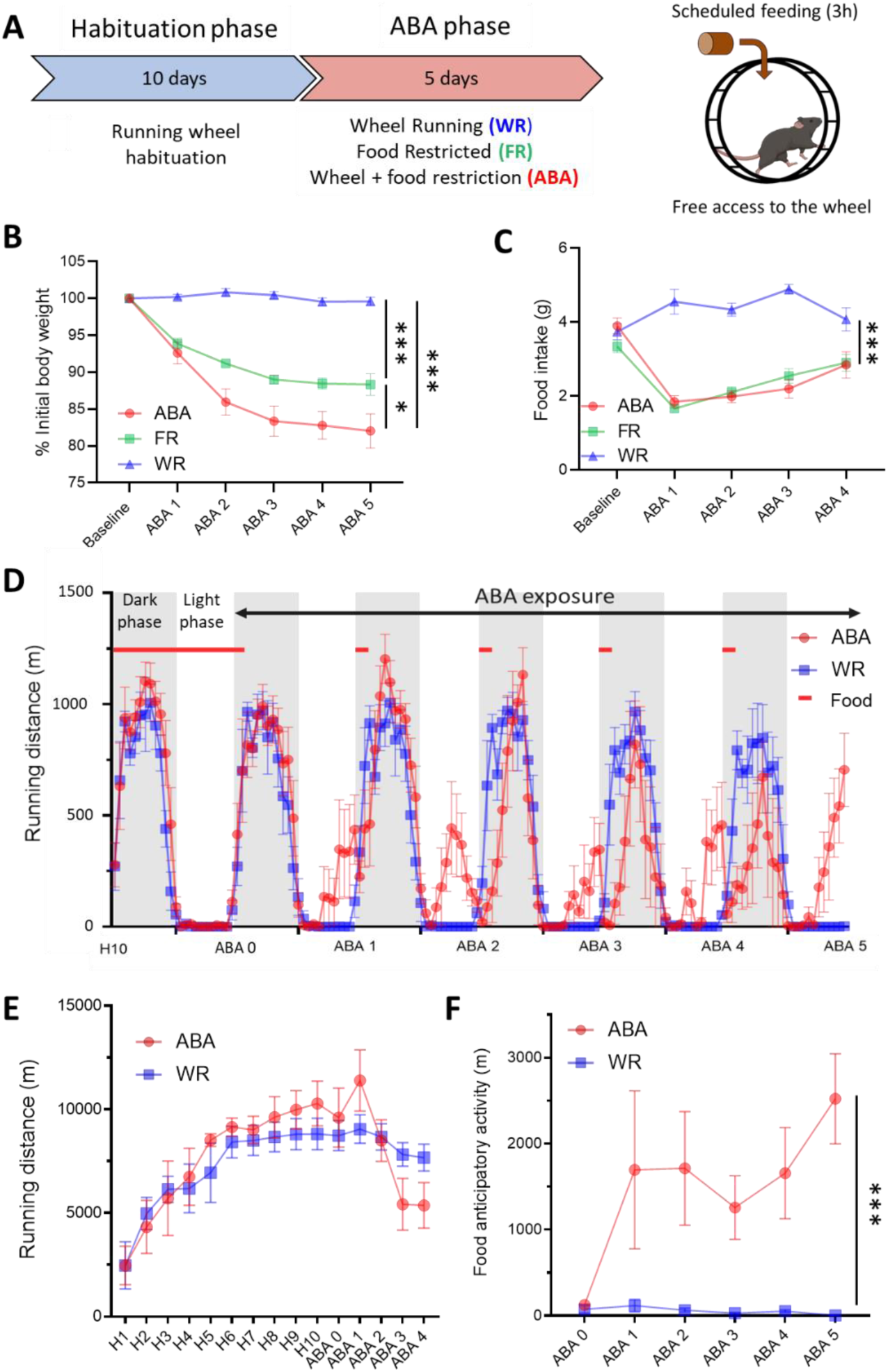
Activity-based anorexia model. (A) Schematic of the ABA experimental design. (B) Average daily body weight variation (C) and food intake during ABA exposure compared to baseline. Control groups are WR and FR animals. (D) Hour by hour distance travelled in the wheel during habituation day 10 (H10) and ABA day 0 to day 5. (E) Overview of daily running wheel activity during the experiment. (F) Food anticipatory activity (FAA) measured during the last 6h of the light phase at H10 and ABA1 to ABA5. Data are means ± SEM. *p<0.05, **p<0.01, ***p<0.001. WR, wheel running control group; FR, food restricted control group; ABA, activity-based anorexia group. See Table S1 for statistical analyses.

### ABA exposure leads to increase VTA_DA_ neuronal activity

We first aimed to determine the impact of exposure to the ABA model on DA neurons in the VTA. Therefore, we performed electrophysiological patch-clamp recordings on *ex vivo* brain slices, allowing measurements of a variety of neuronal electrical properties. The five-days ABA protocol was followed by harvesting the brain just before dark onset, i.e. after FAA occurs in ABA animals and before food exposure for ABA and FR groups. The brain slices were used for cell-attached patch-clamp recordings of VTA_DA_ neurons by using Pitx3-GFP female mice (Fig.2A). ABA exposure did not affect the proportion of spontaneously active VTA_DA_ neurons (Fig.2B), known to display a regular “pace-making” activity [36]. However, the firing frequency of the spontaneously active VTA_DA_ neurons was increased in ABA-exposed animals compared to WR and FR groups (Fig.2C-D and S2B). Thus, ABA-exposed animals display an increased frequency of VTA_DA_ neuronal activity compared to controls.

**Figure 2:**
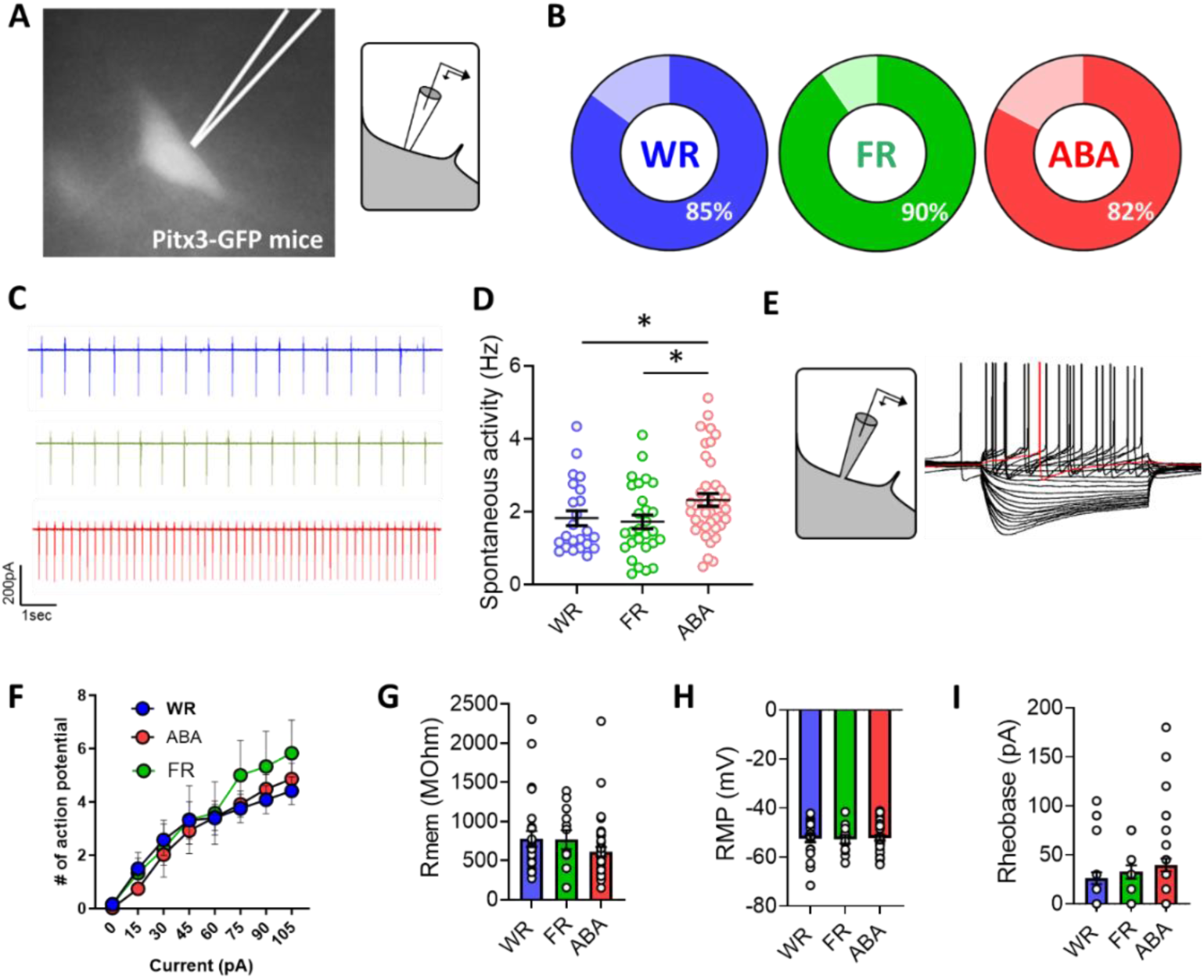
Exposure to the ABA model increase VTADA spontaneous firing frequency but not VTADA intrinsic properties. (A) Representative image of a VTADA neuron of a Pitx3-GFP mouse at high magnification epifluorescence (left) and a schematic representation of the cell-attached patch-clamp mode used (right). (B) Proportion of VTADA neurons found spontaneously active in the 3 experimental groups. (C) Representative traces of the firing pattern measured in VTADA neurons and (D) the associated measured firing rate. (E) Representative traces of voltage responses in VTADA neuron obtained in whole-cell patch-clamp recording. The red trace shows the rehobase. (F) Measure of the frequency of spikes-current curves. (G-I) Measures of membrane resistance (G), resting membrane potential (H), and rheobase (I). Data are means ± SEM. *p<0.05. WR, wheel running control group; FR, food restricted control group; ABA, activity-based anorexia group. See Table S1 for statistical analyses.

We next investigated if this difference in firing frequency was due to changes in cell excitability (i.e. intrinsic electrical properties). Results are presented in Fig.2E-I and Fig.S2C-E where whole-cell patch-clamp recordings were performed allowing to access the intracellular compartment of VTA_DA_ neurons (Fig.2E). Overall, we found that ABA exposure did not affect the intrinsic properties of VTA_DA_ neurons. More precisely, input/output curves, resting membrane potential (RMP), membrane resistance (Rmem), rheobase, action potential (AP) threshold were found unchanged (Fig.2F-I and Fig.S2D-E). To summarize, VTA_DA_ neurons from ABA-exposed animals display an increased firing frequency compared to control groups, which is not due to alterations of basic membrane properties of VTA_DA_ cells.

### Reduced inhibitory transmission from local VTA GABA neurons to VTA_DA_ neurons in ABA-exposed animals

As intrinsic properties of VTA_DA_ cells were unaltered after ABA, the integrity of neuronal inputs they received - influencing neuronal activity by excitation (i.e. glutamatergic transmission) or inhibition (i.e. GABAergic transmission) – was then assessed. We found that the spontaneous glutamatergic excitatory transmission onto VTA_DA_ neurons was unaltered in ABA animals (Fig.S3A-B) whereas spontaneous GABAergic inhibitory transmission was reduced following ABA exposure (Fig.3A-B). Specifically, the frequency of spontaneous inhibitory post-synaptic currents (sIPSC) was decreased in the ABA group whereas the average amplitude of sIPSCs was similar between groups (Fig.3A-B). The difference in IPSC frequency was not found anymore after application of tetrodotoxin (miniature IPSCs, Fig.S3C-D). In ABA-exposed animals, there was a reduction in GABAergic signals onto VTA_DA_ neurons, likely involving a presynaptic but action potential-dependent mechanism. The GABAergic deficits in the ABA condition were specific to the synapse, as we did not observe any alterations of extrasynaptic GABA-A mediated tonic conductances (Fig.S3E-G).

**Figure 3:**
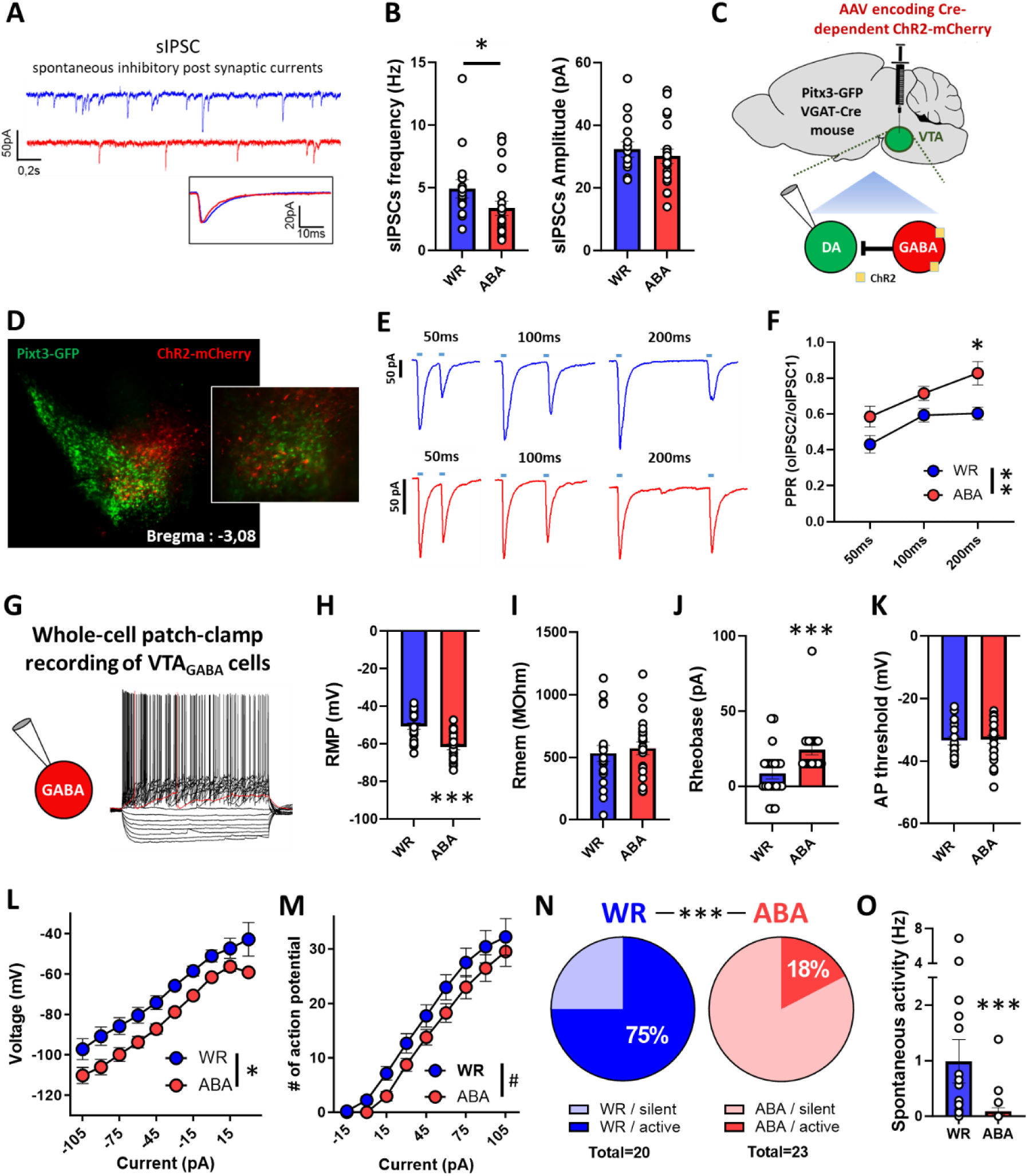
VTAGABA neurons display weaker synapses onto VTADA neurons and decreased excitability in ABA animals. (A) Representative traces of spontaneous IPCSs acquired during voltage clamp recording of VTADA neurons. (B) Measures of sIPSC frequency (left) and amplitude (right). (C) The viral approach to optogenetically target VGAT-expressing neurons in the VTA while patching VTADA neurons. VGAT-Cre; Pitx3-GFP mice were injected with an AAV-ChR2-mCherry into the VTA, allowing the expression of the channel rhodopsin in VGAT-expressing neurons (as shown in D). (E) Example traces of evoked IPSCs with a ≈5mW light stimulation performed in coronal slices. This paired-pulse ratio (PPR) protocol consist in three recordings of two consecutive stimulation with an inter-stimulation interval of 50ms (left), 100ms (middle) and 200ms (right). (F) Measure of the PPR obtained by dividing the amplitude of eIPSC2 with amplitude of eIPSC1 for each time interval. (G) Representative traces of voltage responses in VTAGABA neuron obtained in whole-cell patch-clamp recording. (H-M) Measures of resting membrane potential (H), membrane resistance (I), rheobase (J), action potential threshold (K), current-voltage curves (L) and frequency of spikes-current curves (M). (N) Proportion of VTAGABA neurons found spontaneously active in the 2 experimental groups and (O) the associated measured firing rate. Data are means ± SEM. #p<0.1, *p<0.05, **p<0.01, ***p<0.001. WR, wheel running control group; ABA, activity-based anorexia group. See Table S1 for statistical analyses.

To further assess presynaptic GABA release, a paired-pulse ratio (PPR) protocol was performed consisting of a paired stimulation of a synapse given in a quick succession with 3 different time intervals (Fig.3E and S3H). Electrical stimulation of synapses resulted in an overall increased PPR in the ABA group (FigS3I) which is in accordance with a scenario in which the probability of GABA release is reduced when compared to the WR group [37]. As the electrical stimulation is a synapse non-specific approach, we next asked which input was affected. We predicted that local GABA neurons were involved considering their powerful regulatory function onto dopamine neurons. To investigate this, the PPR was performed using optogenetic stimulations on midbrain GABA neurons surrounding VTA_DA_ cells (Fig.3D-E). This protocol revealed an increased PPR in the ABA group (Fig.3F) showing that local midbrain GABA to VTA_DA_ synapses displayed reduced probability of GABA release. To further corroborate a deficit in local GABAergic VTA interneurons, whole-cell patch-clamp recordings of VTA_GABA_ neurons were performed (Fig.3G). VTA_GABA_ neurons from ABA-exposed animals displayed decreased excitability. More precisely, RMP (Fig.3H) and rheobase (Fig.3J) were found decreased and increased respectively whereas Rmem (Fig.3I) and AP threshold (Fig.3K) were unchanged. This results in a significantly shifted I/V plot (Fig.3L) as well as a trend of a decreased ability to trigger action potentials with increasing steps of current injections (Fig.3M). Confirming this finding, ABA exposure also affects the proportion of spontaneously active VTA_GABA_ neurons (Fig.3N) and therefore reduces their firing frequency compared to WR controls regardless of the chosen experimental unit, the cell (Fig.3O) or the animal (Fig.S3J). Altogether, these data suggest that the inhibitory transmission from local VTA_GABA_ neurons onto VTA_DA_ neurons is weaker in the ABA group compared to the WR group.

### Chemogenetic inhibition of midbrain GABA neurons exacerbates negative energy balance induced by the ABA model while their activation rescues from starvation

To study the causal involvement of ventral midbrain GABA neurons on running wheel activity and food intake, we chemogenetically targeted these neurons. Adeno-associated viruses with Cre-dependent DREADDs (either excitatory hM3Dq or inhibitory hM4Di) were injected in the VTA of vesicular GABA transporter (VGAT)-cre mice, (Fig.4A and S4A-C) allowing DREADD expression in GABA neurons selectively.

**Figure 4:**
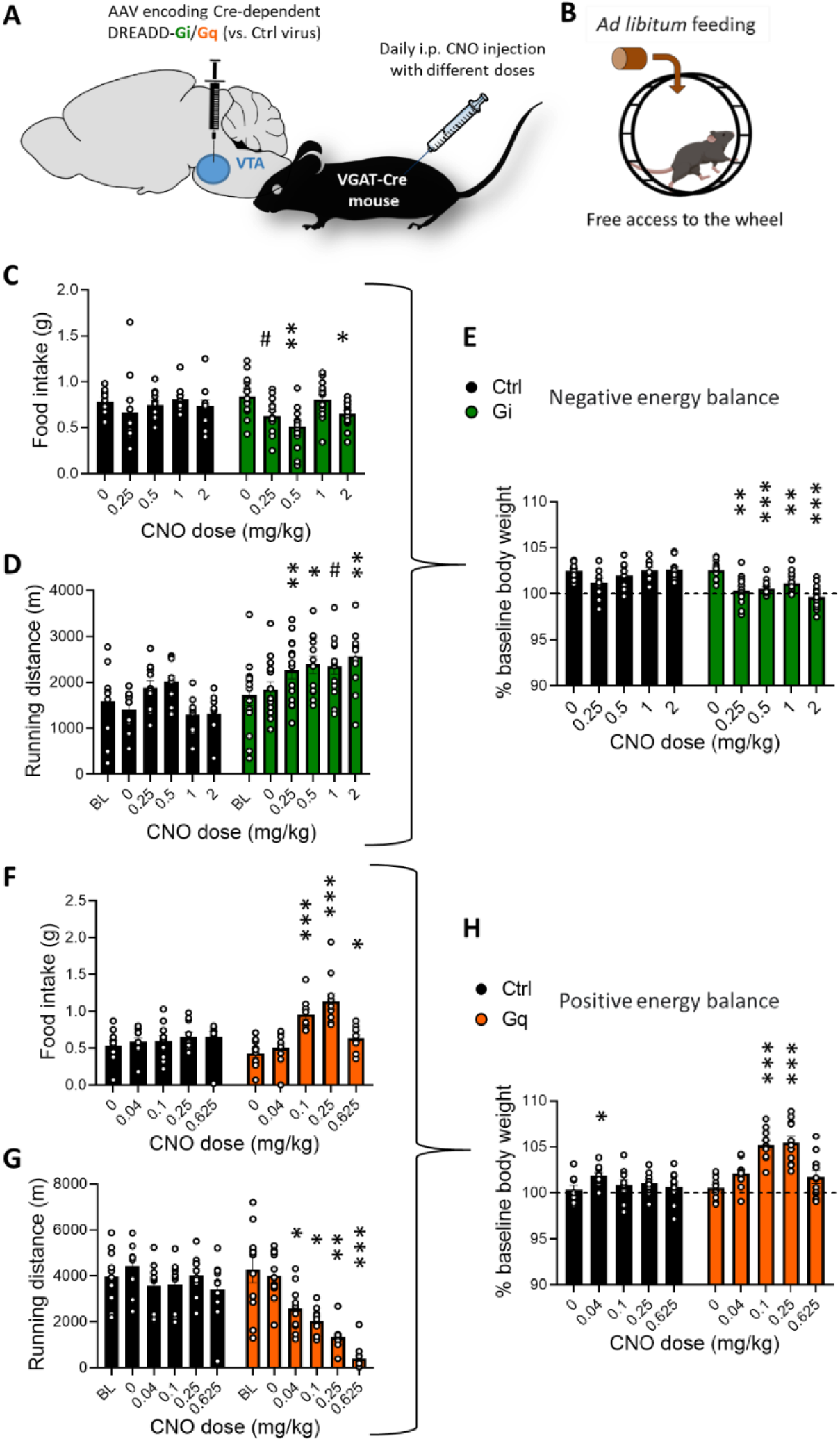
Chemogenetic manipulation of midbrain GABA neurons bidirectionally modulates energy balance in *ad libitum* fed animals. (A) The viral approach to chemogenetically target VGAT-expressing neurons in the midbrain. (B) Schematic of the *ad libitum* feeding and running experimental design. (C-E) Measures in inhibitory DREADD-Gi expressing animals of the dose response to CNO for feeding (C), running (D) and the consequence on body weight variation 3 hours following the injection (E). (F-H) Measures in excitatory DREADD-Gq expressing animals of the dose response to CNO for feeding (F), running (G) and the consequence on body weight variation 3 hours following the injection (H). Data are means ± SEM. #p<0.1, *p<0.05 **p<0.01, ***p<0.001. Gi: DREADD-Gi receptor-expressing mice, Gq: DREADD-Gq receptor-expressing mice, Ctrl: control group, FAA: food anticipatory activity, VTA: ventral tegmental area, VGAT: vesicular GABA transporter, i.p.: intraperitoneal CNO: clozapine-N-Oxide. See Table S1 for statistical analyses.

We first tested whether this GABAergic network regulates food intake and RWA, the main contributors to energy balance, in an *ad libitum* situation (Fig.4B). Different doses of CNO were randomly administered every day. We found that inhibition of midbrain GABA neurons led, for most of the doses used, to decreased food intake (Fig.4C) as well as increased RWA (Fig.4D) within the 3h following CNO administration. This led to a mild but significant decrease in body weight expressed as relative body weight within these 3h and compared to the saline injection day (Fig.4E). In contrast, activating midbrain GABA neurons led to increased food intake and decreased RWA resulting in a significant increase in body weight (Fig.4F-H) when doses of 0.1-0.25mg/kg were used. However, when midbrain GABA neurons are activated at the highest CNO dose tested, running is drastically suppressed and food intake mildly increased suggesting the induction of motor-based side effects. Based on these results, we identified the optimal CNO doses for both chemogenetic inhibition (0.5 mg/kg) and activation (0.25 mg/kg). Importantly, with these doses, we found no alterations in motor coordination nor in DREADD-Gi animals (Fig.S4D) neither in DREADD-Gq animals (Fig.S4E), meaning that the impact on RWA performances was not due to motoric dysfunctions. Moreover, it is important to note that CNO had no effect in control animals that express the control fluorophore, regardless of the tested dose. To summarize, midbrain GABA neurons are involved in energy balance regulation by acting on both feeding and exercise in an *ad libitum* situation.

We next studied the involvement of ventral midbrain GABA neurons on RWA and food intake in ABA-exposed animals (Fig.5A-B). As shown in Fig.5C-F, animals expressing the inhibitory DREADD-Gi and the control group were exposed to ABA and injected with 0.5 mg/kg CNO every day 5h before food exposure (when most of the animals start expressing FAA). Inhibiting midbrain GABA neurons increased the ABA-induced body weight loss (Fig.5C), compared to ABA-controls. There were no changes in daily food intake or total RWA (Fig.5D and S5C). However, light phase RWA and FAA were increased in the DREADD-Gi group (Fig5E and S5A). Thus, inhibition of midbrain GABA neurons accelerated the development of FAA leading to an increased vulnerability to the ABA model (Fig.5C and 5F).

**Figure 5:**
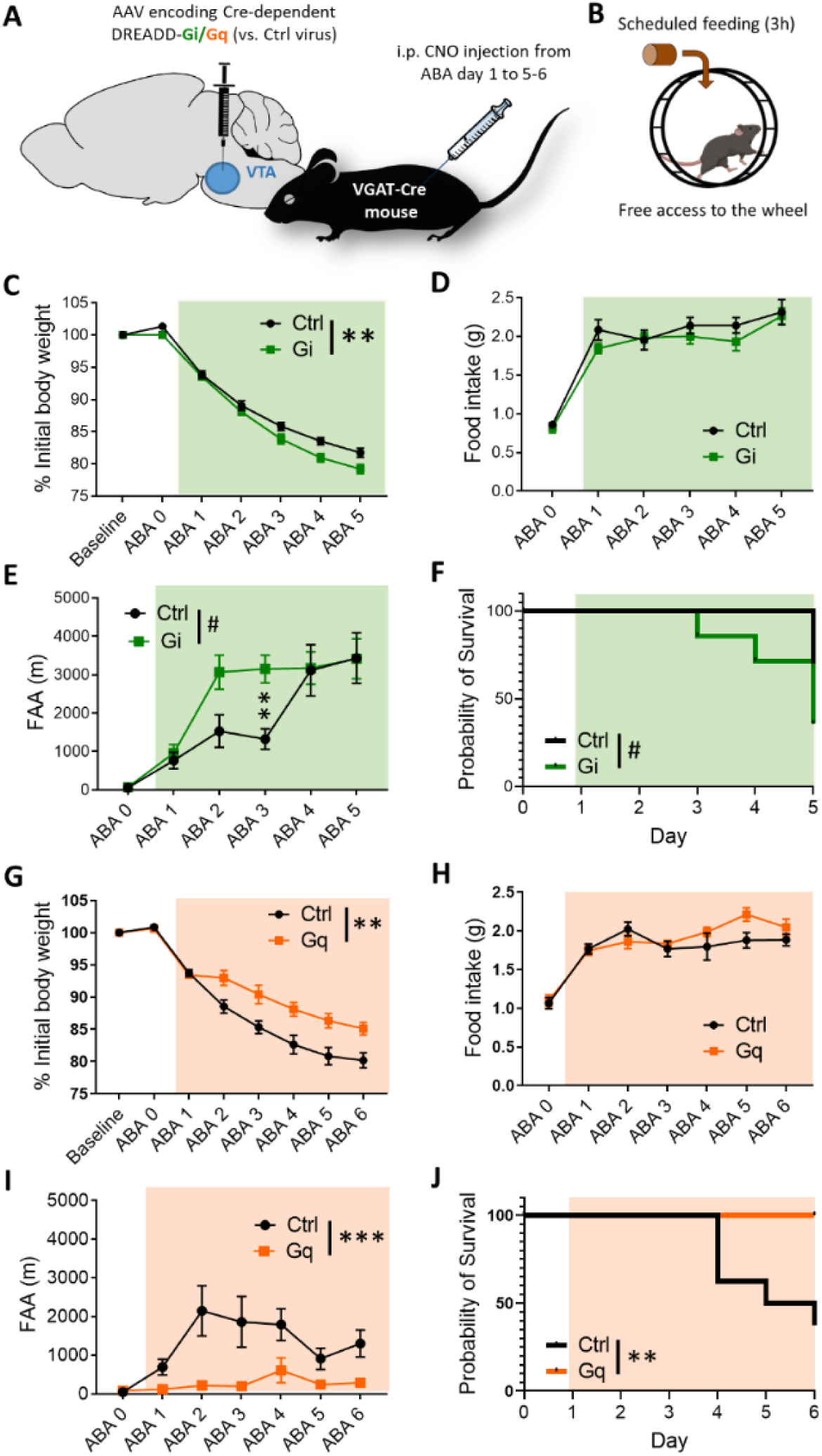
Chemogenetic inhibition of midbrain GABA neurons exacerbates negative energy balance induced by the ABA model while their activation partially rescues it. (A) The viral approach to chemogenetically target VGAT-expressing neurons in the midbrain, VGAT-Cre mice were injected with AAV-hM4D(Gi)-mCherry or AAV-hM3D(Gq)-mCherry into the VTA, allowing the expression of the DREADD-Gi or DREADD-Gq receptor respectively. CNO, via its action on the DREADD-Gq receptor, inhibit (Gi) or activate (Gq) these neurons. (B) Schematic of the ABA experimental design. (C-F) DREADD-Gi expressing animals daily injected with 0.5 mg/kg CNO 5h before feeding. Measures over the 5 days of ABA of body weight loss (C), food intake (D), FAA measured as running distance travelled 5h before dark onset (beginning of food exposure, E) and ‘‘survival’’ measured as the percentage of animals above 80% of initial body weight (F). (G-J) DREADD-Gq expressing animals daily injected (twice a day: 5h before feeding and after the 3h feeding period) with 0.25 mg/kg CNO. Measures over the 6 days of ABA of body weight loss (G), food intake (H), FAA measured as running distance travelled 5h before dark onset (beginning of food exposure, I) and “survival’’ measured as the percentage of animals above 80% of initial body weight (J). Data are means ± SEM. #p<0.1, **p<0.01, ***p<0.001. Gi: DREADD-Gi receptor-expressing mice, Gq: DREADD-Gq receptor-expressing mice, Ctrl: control group, FAA: food anticipatory activity, VTA: ventral tegmental area, VGAT: vesicular GABA transporter, i.p.: intraperitoneal CNO: clozapine-N-Oxide. See Table S1 for statistical analyses.

In Fig.5G-J, the opposite experiment was performed with activating DREADD-Gq receptors. We found that activation of midbrain GABA neurons with 0.25 mg/kg CNO twice a day was protective against body weight loss (Fig.5G). Daily measurements of food intake or total RWA did not reveal any differences (Fig.5H and S5D) but FAA was drastically suppressed (Fig.5I and S5B). These data suggest that excessive running that develops along ABA exposure can be suppressed by increasing midbrain GABA neuronal activity, resulting in a better survival to the ABA model (Fig.5G and J). Altogether, these data suggest that restoring the excitability of midbrain GABA neurons via chemogenetic activation prevented their attrition in the task, i.e. reducing body weight loss by decreasing running wheel activity. Conversely, chemogenetic inhibition increased vulnerability to the ABA model by increasing FAA.

## DISCUSSION

We here identified a VTA neural circuit that underlies hyperactivity and excessive weight loss in an animal model for anorexia nervosa. ABA-exposed mice display increased activity of VTA_DA_ neurons co-occurring with a decreased GABA release probability and reduced neuronal excitability of local VTA_GABA_ cells, the second likely responsible for the first. These results support that the midbrain dopamine system undergoes plasticity under the condition of restricted feeding and voluntary exercise. Restoring activity of GABA neurons by chemogenetics decreases vulnerability to the ABA model by suppression of excessive running and reduction of body weight loss. We identified a novel mechanism explaining how negative energy balance leads to dysregulation of the midbrain dopamine system that in turn reinforces maladaptive behaviors such as hyperactivity.

Our results suggest that disinhibition of the dopamine system may be a key candidate for the establishment of negative energy balance. There are little clinical data on GABA (dys)functions in AN and GABAergic drugs have hardly been studied in AN therapy [38, 39]. A few preclinical studies investigated the link between central alterations of GABA transmission and AN-like symptoms such as hyperactivity, decreased food intake and anxiety levels in AN rodent models [40]. Decreased GABA levels were found in the hypothalamus of ABA rats and indirect rescue of GABA levels (with the neuropeptide kisspeptin) increased food intake [41]. Within amygdala circuits, ABA-exposure compromised inhibition [42]. In another study, increased inhibition in rodent hippocampal pyramidal neurons was found protective against the ABA-induced weight loss by indirectly suppressing hyperactivity [42]. Therefore, Aoki and colleagues proposed that ABA vulnerability partly relies on GABAergic alterations and increased GABA transmission may rescue maladaptive behaviors such as excessive exercise [30]. To our knowledge, the present study is the first to provide evidence for reduced VTA_GABA_ transmission being involved in AN-like symptoms, such as hyperactivity, likely due to increased VTA_DA_ activity.

One limitation of this study is that we cannot exclude that other sources of GABA than those from the VTA contribute to the alterations we found in VTA_DA_ activity. Although the VTA was chemogenetically transduced, there was infection of GABA cells near to the VTA in some animals. However, DREADDs expression outside the VTA varied from one individual to another (in location and size) but DREADD receptors were consistently expressed within the VTA in all animals. Hence, considering our electrophysiological data, we concluded that decreased excitability of VTA_GABA_ neurons is a major actor to induce hyperactivity in ABA-exposed animals. Furthermore, there are also other GABAergic inputs to VTA dopamine neurons from outside of the midbrain that may be affected by ABA exposures, such as from the nucleus accumbens [43], and lateral hypothalamus [34]. Whether such inputs are also altered remains to be addressed.

Additionally, further studies will be necessary to identify the molecular mechanisms behind GABAergic alterations. Recent case reports showed that short term treatment of AN patients with leptin, improved their mood and reduced hyperactivity [44, 45]. As mood and hyperlocomotion involve dopamine transmission and leptin is known to modulate the VTA, including VTA_GABA_ neurons [46, 47] and the drive for exercise [48–50], decreased leptin levels may be a causal factor in the reduced GABAergic signaling we found.

The results obtained in this study are in accordance with the hypothesized relationship between the hyperdopaminergic function and the regulation of energy expenditure [51]: low dopamine function leads to restrictive behaviors (conserve energy/exploit environment) whereas higher dopamine function drives expansive behaviors (expend energy/explore environment). This is supported by a growing number of studies manipulating dopamine transmission in ABA-exposed animals and showing that increased dopamine transmission leads to higher vulnerability to body weight loss [22, 26, 29, 52]. Additionally, running wheel activity has been shown to increase striatal DA release and induce a hyperdopaminergic state [53, 54]. Chemogenetic activation of VTA_DA_ neurons and neurons projecting from the VTA to the NAc leads to hyperlocomotion [17], further supporting the idea that activation of the DA reward system and hyperactivity are causally linked. Surprisingly, one study found that activating VTA to NAc neurons prevented and rescued body weight loss (by increasing food intake) induced by the ABA model [55], but in that study non-dopaminergic projection neurons were likely also targeted. Recently, a biphasic model of AN has been proposed with 1) a hyperdopaminergic state in the acute phase, accompanied by symptoms such as hyperactivity and 2) a chronic low dopamine function resulting in behavioral rigidity (Beeler and Burghardt, 2023). This model supports the clinical observations of low DA metabolite levels and increased D2 binding in recovered AN patients. That said, the mechanisms and brain circuits linking AN-like symptoms and the dopamine system are still poorly understood. In this context, our data suggest that reduced VTA_GABA_ neuron activity is causal to negative energy balance by increasing dopamine function leading to the development of maladaptive behaviors such as hyperactivity.

The ABA paradigm is a well-established rodent model of AN [23, 56] and the development of FAA is a hallmark and intriguing phenomenon. We hypothesized that FAA, and more generally hyperlocomotion, could occur because of an urge to exercise originating from alterations of VTA neuronal activity. Indeed, in patients with AN the hyperactivity that develops despite negative consequences may originate from alterations at the motivational level [3, 57] and future research would be needed to study the competition between motivation to eat and motivation to exercise that the ABA model cannot fully disentangle. Moreover, increasing dopamine function does not only underlie hyperactivity but also reward-related decision making [58] that resemble the insensitivity to losses observed in AN patients [59]. Therefore, studying reward processing in more depth would be of great interest in the field. Nonetheless, this study suggests that neural circuits in the VTA, known to be involved in other forms of compulsive/addictive behaviors [60], may participate to the development of excessive exercise in eating disorders.

Altogether, our data brings new insights on the impact of running and food restriction on the reward system, allowing to better understand the neuronal mechanisms involved in energy balance regulation. Interestingly, Beeler and Burghardt recently hypothesized that the combination of caloric restriction with exercise, triggers an escalating spiral of increasing dopamine reinforcing weight-loss behaviors [25]. Our results support this hypothesis. The results found in this study also suggest for the first time that a discrete neuronal population, namely VTA_GABA_ neurons, is directly contributing to hyperactivity and favor restlessness. Indeed, ABA-exposed mice expend more energy because they may be “restless” and activation of VTA_GABA_ neurons decrease excessive locomotor activity. This is particularly relevant in the context of AN since restlessness is a symptom of AN [4, 61] and our results may contribute to a new therapeutic avenue to explore. Overall this study contributes to further understanding of how changes in VTA neuronal activity and neural circuit interact to impact on the susceptibility to develop anorectic behavior in the ABA model.

## ACKNOWLEDGEMENTS

We would like to thank the entire Adan and Meye Labs for discussions and critical reading of the manuscript. This research was supported by the Dutch Research Council NWO ALWOP.137, OCENW.M.22.111 and Gravitation grant 024.004.012; to RAH and the Fyssen Fundation; to FD.

## AUTHOR CONTRIBUTIONS

R.A. and F.D. designed experiments with the contribution of F.M. F.D. performed behavioral, chemogenetic and *ex vivo* electrophysiology experiments. E.B., L.D., J.H and M.V. helped with behavioral experiments. F.D., I.W.D., M.L and K.K. performed stereotactic surgeries. F.D. performed analysis on all experiments. F.D., R.A. and F.M. wrote the manuscript with input from all authors.

## SUPPLEMENTARY METHOD DETAILS

### ROTAROD

A ROTA-ROD treadmill apparatus (Cat.No.47600, UGO BASILE) was used to assess motor coordination and balance of mice under CNO. The protocol consisted of 2 days (with one “experiment-free” day in between) allowing a randomized saline/CNO comparison were all animals are submitted to the ROTAROD 3 times each day, starting 20min after injection and with 15min in between every ROTAROD session. One session consisted of 300 sec starting with a 4 rotation per minute (RPM) to a 60 RPM (+1 RPM every 5 seconds) and the latency to fall was recorded. All mice had a training session 15min prior the first session on experimental day 1. The average of the 3 measurements was plotted for both saline and CNO conditions.

### Whole-Cell Patch Clamp Recording

Spontaneous excitatory post-synaptic currents (EPSCs) we recorded with neurons clamped at -65mV). Tetrodotoxin (TTX, 1µM, Tocris 1078) was added to the bath in order to isolate miniature inhibitory postsynaptic currents (mIPSC) and Picrotoxin (PTX, 100µM, Tocris) was used in order to measure GABA_A_ receptors-mediated tonic inhibition and neurons were voltage clamped at -70mV. Recordings of sIPSCs, and mIPSCs were performed at least 10min after patch for 1min. Tonic GABAergic conductance was defined as a difference between the baseline holding current and that obtained after blocking for 5 minutes GABA_A_ receptors with PTX. Electrical stimulations were performed with a bipolar concentric stimulating electrode using a stimulator (Isoflex, AMPI) with electric pulses of 0.18ms; 50-3000μA.

### Immunohistochemistry

The viral injection sites were confirmed by immunohistochemistry and fluorescent microscopy and all operated animals were included in the study. For the chemogenetic experiments, mice were transcardially perfused with ice-cold 4% paraformaldehyde in PBS under anesthesia after behavioral experiments. Brains were harvested, postfixed overnight and washed in PBS. 50μm coronal sections were obtained using a vibratome VT1000s (Leica) and incubated in a blocking solution (5% goat serum, 2.5% BSA, 0.2% Tritonx100 in PBS, pH 7.4) for 1h at room temperature (RT). Slices were labeled overnight at 4°C with primary antibodies against TH (mouse; 1:1000, MAB318, Merck). Sections were incubated with fluorescent secondary antibodies (goat anti mouse 488) for 2h at RT and then mounted on slides and coverslipped with the fluorsave reagent (FluorSave Reagent, Merk Millipore). Digital images of colocalization of TH immunostaining and mCherry (AAV-DREADD-mCherry) were acquired using an epifluorescent microscope (Olympus). For the optogenetic experiments, the slices of 250μm were postfixed right after patch-clamp experiment to similarly check for colocalization of endogenous GFP (Pitx3-GFP mice) and mCherry (AAV-ChR2-mCherry).

**Table S1.**
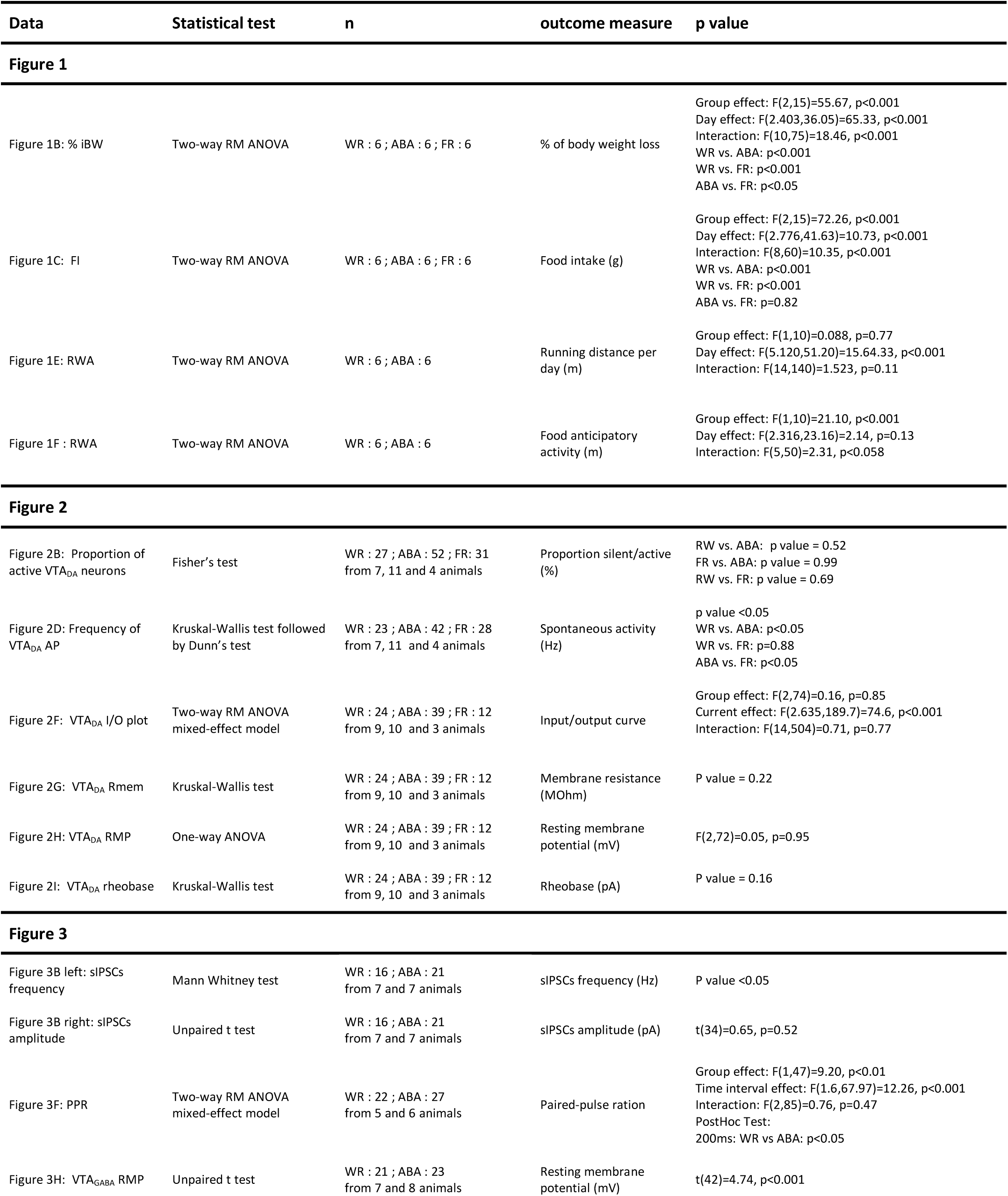
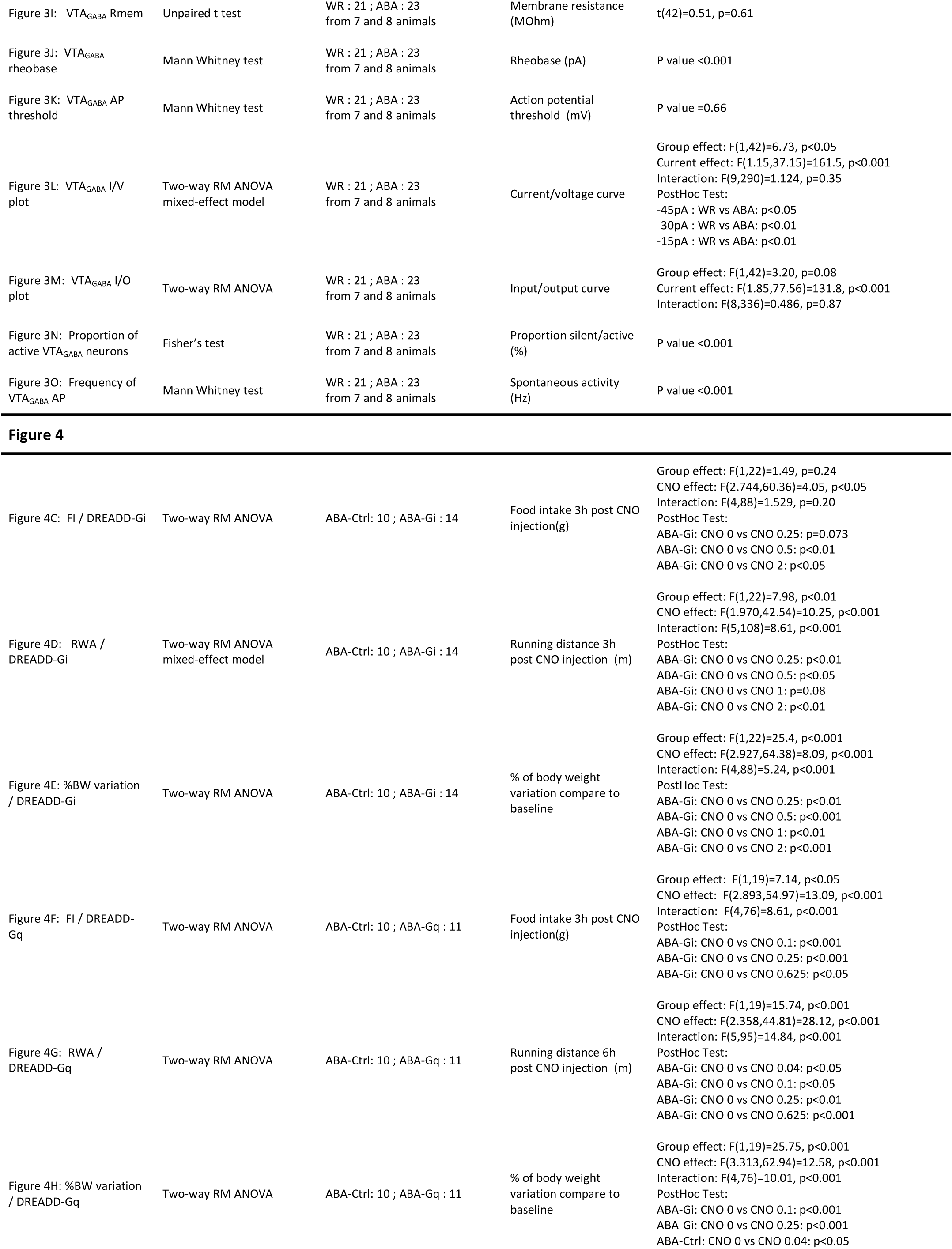
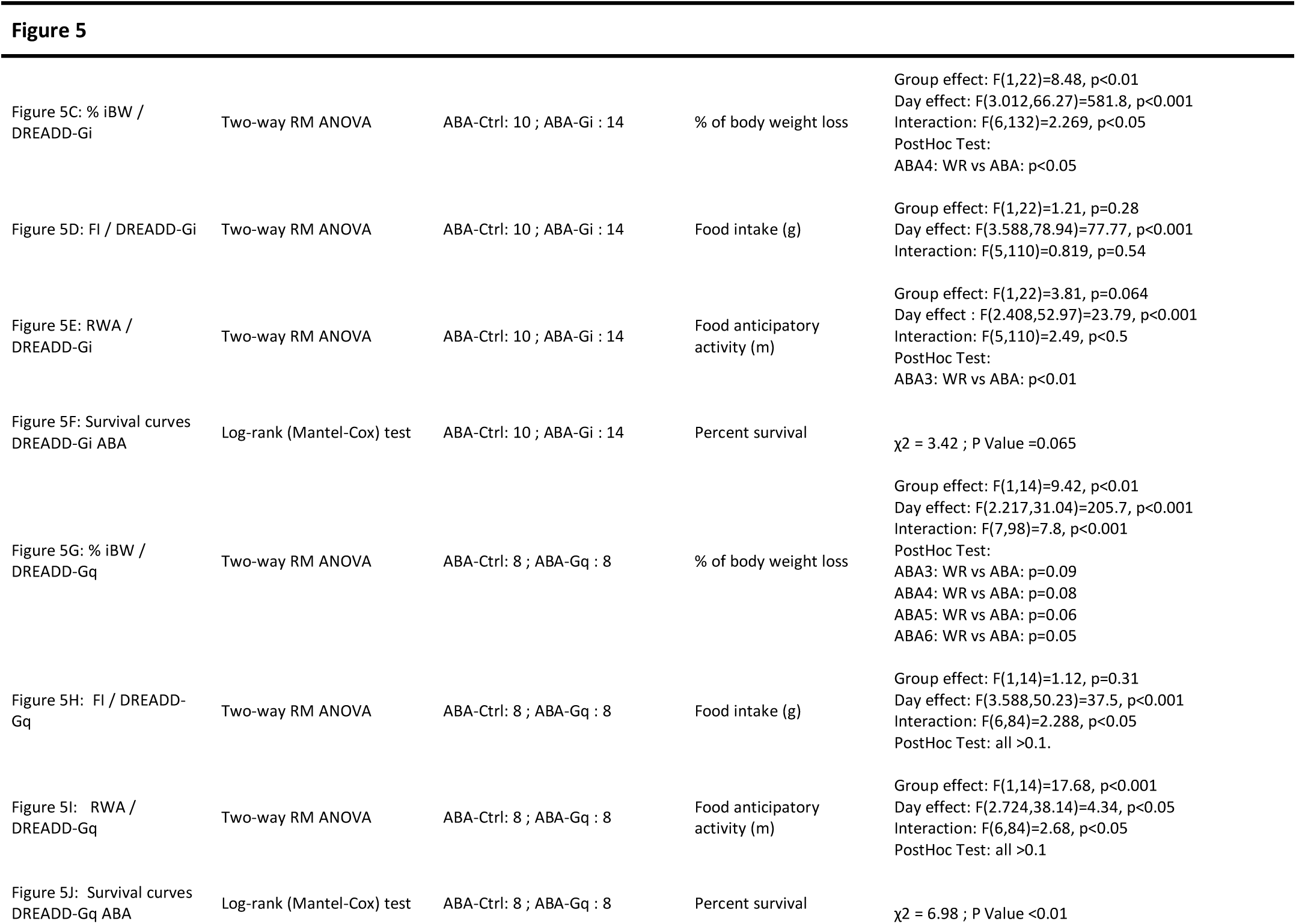
Summary of statistical analysis

| Data | Statistical test | n | outcome measure | p value |
| --- | --- | --- | --- | --- |
| **Figure 1** |  |  |  |  |
| Figure 1B: % iBW | Two-way RM ANOVA | WR : 6 ; ABA : 6 ; FR : 6 | % of body weight loss | Group effect: $F(2,15)=55.67$, $p<0.001$<br>Day effect: $F(2.403,36.05)=65.33$, $p<0.001$<br>Interaction: $F(10,75)=18.46$, $p<0.001$<br>WR vs. ABA: $p<0.001$<br>WR vs. FR: $p<0.001$<br>ABA vs. FR: $p<0.05$ |
| Figure 1C: FI | Two-way RM ANOVA | WR : 6 ; ABA : 6 ; FR : 6 | Food intake (g) | Group effect: $F(2,15)=72.26$, $p<0.001$<br>Day effect: $F(2.776,41.63)=10.73$, $p<0.001$<br>Interaction: $F(8,60)=10.35$, $p<0.001$<br>WR vs. ABA: $p<0.001$<br>WR vs. FR: $p<0.001$<br>ABA vs. FR: $p=0.82$ |
| Figure 1E: RWA | Two-way RM ANOVA | WR : 6 ; ABA : 6 | Running distance per day (m) | Group effect: $F(1,10)=0.088$, $p=0.77$<br>Day effect: $F(5.120,51.20)=15.64.33$, $p<0.001$<br>Interaction: $F(14,140)=1.523$, $p=0.11$ |
| Figure 1F: RWA | Two-way RM ANOVA | WR : 6 ; ABA : 6 | Food anticipatory activity (m) | Group effect: $F(1,10)=21.10$, $p<0.001$<br>Day effect: $F(2.316,23.16)=2.14$, $p=0.13$<br>Interaction: $F(5,50)=2.31$, $p<0.058$ |
| **Figure 2** |  |  |  |  |
| Figure 2B: Proportion of active $VTA_{DA}$ neurons | Fisher's test | WR : 27 ; ABA : 52 ; FR: 31 from 7, 11 and 4 animals | Proportion silent/active (%) | RW vs. ABA: p value = 0.52<br>FR vs. ABA: p value = 0.99<br>RW vs. FR: p value = 0.69 |
| Figure 2D: Frequency of $VTA_{DA}$ AP | Kruskal-Wallis test followed by Dunn's test | WR : 23 ; ABA : 42 ; FR : 28 from 7, 11 and 4 animals | Spontaneous activity (Hz) | p value <0.05<br>WR vs. ABA: $p<0.05$<br>WR vs. FR: $p=0.88$<br>ABA vs. FR: $p<0.05$ |
| Figure 2F: $VTA_{DA}$ I/O plot | Two-way RM ANOVA mixed-effect model | WR : 24 ; ABA : 39 ; FR : 12 from 9, 10 and 3 animals | Input/output curve | Group effect: $F(2,74)=0.16$, $p=0.85$<br>Current effect: $F(2.635,189.7)=74.6$, $p<0.001$<br>Interaction: $F(14,504)=0.71$, $p=0.77$ |
| Figure 2G: $VTA_{DA}$ Rmem | Kruskal-Wallis test | WR : 24 ; ABA : 39 ; FR : 12 from 9, 10 and 3 animals | Membrane resistance (MΩm) | P value = 0.22 |
| Figure 2H: $VTA_{DA}$ RMP | One-way ANOVA | WR : 24 ; ABA : 39 ; FR : 12 from 9, 10 and 3 animals | Resting membrane potential (mV) | $F(2,72)=0.05$, $p=0.95$ |
| Figure 2I: $VTA_{DA}$ rheobase | Kruskal-Wallis test | WR : 24 ; ABA : 39 ; FR : 12 from 9, 10 and 3 animals | Rheobase (pA) | P value = 0.16 |
| **Figure 3** |  |  |  |  |
| Figure 3B left: sIPSCs frequency | Mann Whitney test | WR : 16 ; ABA : 21 from 7 and 7 animals | sIPSCs frequency (Hz) | P value <0.05 |
| Figure 3B right: sIPSCs amplitude | Unpaired t test | WR : 16 ; ABA : 21 from 7 and 7 animals | sIPSCs amplitude (pA) | $t(34)=0.65$, $p=0.52$ |
| Figure 3F: PPR | Two-way RM ANOVA mixed-effect model | WR : 22 ; ABA : 27 from 5 and 6 animals | Paired-pulse ration | Group effect: $F(1,47)=9.20$, $p<0.01$<br>Time interval effect: $F(1.6,67.97)=12.26$, $p<0.001$<br>Interaction: $F(2,85)=0.76$, $p=0.47$<br>PostHoc Test:<br>200ms: WR vs ABA: $p<0.05$ |
| Figure 3H: $VTA_{GABA}$ RMP | Unpaired t test | WR : 21 ; ABA : 23 from 7 and 8 animals | Resting membrane potential (mV) | $t(42)=4.74$, $p<0.001$ |
| Figure 3I: VTA<sub>GABA</sub> Rmem | Unpaired t test | WR : 21 ; ABA : 23<br>from 7 and 8 animals | Membrane resistance<br>(MΩm) | t(42)=0.51, p=0.61 |
| Figure 3J: VTA<sub>GABA</sub><br>rheobase | Mann Whitney test | WR : 21 ; ABA : 23<br>from 7 and 8 animals | Rheobase (pA) | P value <0.001 |
| Figure 3K: VTA<sub>GABA</sub> AP<br>threshold | Mann Whitney test | WR : 21 ; ABA : 23<br>from 7 and 8 animals | Action potential<br>threshold (mV) | P value =0.66 |
| Figure 3L: VTA<sub>GABA</sub> I/V<br>plot | Two-way RM ANOVA<br>mixed-effect model | WR : 21 ; ABA : 23<br>from 7 and 8 animals | Current/voltage curve | Group effect: F(1,42)=6.73, p<0.05<br>Current effect: F(1.15,37.15)=161.5, p<0.001<br>Interaction: F(9,290)=1.124, p=0.35<br>PostHoc Test:<br>-45pA : WR vs ABA: p<0.05<br>-30pA : WR vs ABA: p<0.01<br>-15pA : WR vs ABA: p<0.01 |
| Figure 3M: VTA<sub>GABA</sub> I/O<br>plot | Two-way RM ANOVA | WR : 21 ; ABA : 23<br>from 7 and 8 animals | Input/output curve | Group effect: F(1,42)=3.20, p=0.08<br>Current effect: F(1.85,77.56)=131.8, p<0.001<br>Interaction: F(8,336)=0.486, p=0.87 |
| Figure 3N: Proportion of<br>active VTA<sub>GABA</sub> neurons | Fisher's test | WR : 21 ; ABA : 23<br>from 7 and 8 animals | Proportion silent/active<br>(%) | P value <0.001 |
| Figure 3O: Frequency of<br>VTA<sub>GABA</sub> AP | Mann Whitney test | WR : 21 ; ABA : 23<br>from 7 and 8 animals | Spontaneous activity<br>(Hz) | P value <0.001 |
| **Figure 4** |  |  |  |  |
| Figure 4C: FI / DREADD-Gi | Two-way RM ANOVA | ABA-Ctrl: 10 ; ABA-Gi : 14 | Food intake 3h post CNO<br>injection(g) | Group effect: F(1,22)=1.49, p=0.24<br>CNO effect: F(2.744,60.36)=4.05, p<0.05<br>Interaction: F(4,88)=1.529, p=0.20<br>PostHoc Test:<br>ABA-Gi: CNO 0 vs CNO 0.25: p=0.073<br>ABA-Gi: CNO 0 vs CNO 0.5: p<0.01<br>ABA-Gi: CNO 0 vs CNO 2: p<0.05 |
| Figure 4D: RWA /<br>DREADD-Gi | Two-way RM ANOVA<br>mixed-effect model | ABA-Ctrl: 10 ; ABA-Gi : 14 | Running distance 3h<br>post CNO injection (m) | Group effect: F(1,22)=7.98, p<0.01<br>CNO effect: F(1.970,42.54)=10.25, p<0.001<br>Interaction: F(5,108)=8.61, p<0.001<br>PostHoc Test:<br>ABA-Gi: CNO 0 vs CNO 0.25: p<0.01<br>ABA-Gi: CNO 0 vs CNO 0.5: p<0.05<br>ABA-Gi: CNO 0 vs CNO 1: p=0.08<br>ABA-Gi: CNO 0 vs CNO 2: p<0.01 |
| Figure 4E: %BW variation<br>/ DREADD-Gi | Two-way RM ANOVA | ABA-Ctrl: 10 ; ABA-Gi : 14 | % of body weight<br>variation compare to<br>baseline | Group effect: F(1,22)=25.4, p<0.001<br>CNO effect: F(2.927,64.38)=8.09, p<0.001<br>Interaction: F(4,88)=5.24, p<0.001<br>PostHoc Test:<br>ABA-Gi: CNO 0 vs CNO 0.25: p<0.01<br>ABA-Gi: CNO 0 vs CNO 0.5: p<0.001<br>ABA-Gi: CNO 0 vs CNO 1: p<0.01<br>ABA-Gi: CNO 0 vs CNO 2: p<0.001 |
| Figure 4F: FI / DREADD-Gq | Two-way RM ANOVA | ABA-Ctrl: 10 ; ABA-Gq : 11 | Food intake 3h post CNO<br>injection(g) | Group effect: F(1,19)=7.14, p<0.05<br>CNO effect: F(2.893,54.97)=13.09, p<0.001<br>Interaction: F(4,76)=8.61, p<0.001<br>PostHoc Test:<br>ABA-Gi: CNO 0 vs CNO 0.1: p<0.001<br>ABA-Gi: CNO 0 vs CNO 0.25: p<0.001<br>ABA-Gi: CNO 0 vs CNO 0.625: p<0.05 |
| Figure 4G: RWA /<br>DREADD-Gq | Two-way RM ANOVA | ABA-Ctrl: 10 ; ABA-Gq : 11 | Running distance 6h<br>post CNO injection (m) | Group effect: F(1,19)=15.74, p<0.001<br>CNO effect: F(2.358,44.81)=28.12, p<0.001<br>Interaction: F(5,95)=14.84, p<0.001<br>PostHoc Test:<br>ABA-Gi: CNO 0 vs CNO 0.04: p<0.05<br>ABA-Gi: CNO 0 vs CNO 0.1: p<0.05<br>ABA-Gi: CNO 0 vs CNO 0.25: p<0.01<br>ABA-Gi: CNO 0 vs CNO 0.625: p<0.001 |
| Figure 4H: %BW variation<br>/ DREADD-Gq | Two-way RM ANOVA | ABA-Ctrl: 10 ; ABA-Gq : 11 | % of body weight<br>variation compare to<br>baseline | Group effect: F(1,19)=25.75, p<0.001<br>CNO effect: F(3.313,62.94)=12.58, p<0.001<br>Interaction: F(4,76)=10.01, p<0.001<br>PostHoc Test:<br>ABA-Gi: CNO 0 vs CNO 0.1: p<0.001<br>ABA-Gi: CNO 0 vs CNO 0.25: p<0.001<br>ABA-Ctrl: CNO 0 vs CNO 0.04: p<0.05 |
| **Figure 5** |  |  |  |  |
| Figure 5C: % iBW / DREADD-Gi | Two-way RM ANOVA | ABA-Ctrl: 10 ; ABA-Gi : 14 | % of body weight loss | Group effect: $F(1,22)=8.48$, $p<0.01$<br>Day effect: $F(3.012,66.27)=581.8$, $p<0.001$<br>Interaction: $F(6,132)=2.269$, $p<0.05$<br>PostHoc Test:<br>ABA4: WR vs ABA: $p<0.05$ |
| Figure 5D: FI / DREADD-Gi | Two-way RM ANOVA | ABA-Ctrl: 10 ; ABA-Gi : 14 | Food intake (g) | Group effect: $F(1,22)=1.21$, $p=0.28$<br>Day effect: $F(3.588,78.94)=77.77$, $p<0.001$<br>Interaction: $F(5,110)=0.819$, $p=0.54$ |
| Figure 5E: RWA / DREADD-Gi | Two-way RM ANOVA | ABA-Ctrl: 10 ; ABA-Gi : 14 | Food anticipatory activity (m) | Group effect: $F(1,22)=3.81$, $p=0.064$<br>Day effect : $F(2.408,52.97)=23.79$, $p<0.001$<br>Interaction: $F(5,110)=2.49$, $p<0.5$<br>PostHoc Test:<br>ABA3: WR vs ABA: $p<0.01$ |
| Figure 5F: Survival curves DREADD-Gi ABA | Log-rank (Mantel-Cox) test | ABA-Ctrl: 10 ; ABA-Gi : 14 | Percent survival | $\chi^2 = 3.42$ ; P Value =0.065 |
| Figure 5G: % iBW / DREADD-Gq | Two-way RM ANOVA | ABA-Ctrl: 8 ; ABA-Gq : 8 | % of body weight loss | Group effect: $F(1,14)=9.42$, $p<0.01$<br>Day effect: $F(2.217,31.04)=205.7$, $p<0.001$<br>Interaction: $F(7,98)=7.8$, $p<0.001$<br>PostHoc Test:<br>ABA3: WR vs ABA: $p=0.09$<br>ABA4: WR vs ABA: $p=0.08$<br>ABA5: WR vs ABA: $p=0.06$<br>ABA6: WR vs ABA: $p=0.05$ |
| Figure 5H: FI / DREADD-Gq | Two-way RM ANOVA | ABA-Ctrl: 8 ; ABA-Gq : 8 | Food intake (g) | Group effect: $F(1,14)=1.12$, $p=0.31$<br>Day effect: $F(3.588,50.23)=37.5$, $p<0.001$<br>Interaction: $F(6,84)=2.288$, $p<0.05$<br>PostHoc Test: all $>0.1$. |
| Figure 5I: RWA / DREADD-Gq | Two-way RM ANOVA | ABA-Ctrl: 8 ; ABA-Gq : 8 | Food anticipatory activity (m) | Group effect: $F(1,14)=17.68$, $p<0.001$<br>Day effect: $F(2.724,38.14)=4.34$, $p<0.05$<br>Interaction: $F(6,84)=2.68$, $p<0.05$<br>PostHoc Test: all $>0.1$ |
| Figure 5J: Survival curves DREADD-Gq ABA | Log-rank (Mantel-Cox) test | ABA-Ctrl: 8 ; ABA-Gq : 8 | Percent survival | $\chi^2 = 6.98$ ; P Value $<0.01$ |

**Supplementary Figure 1. (related to Figure 1).**
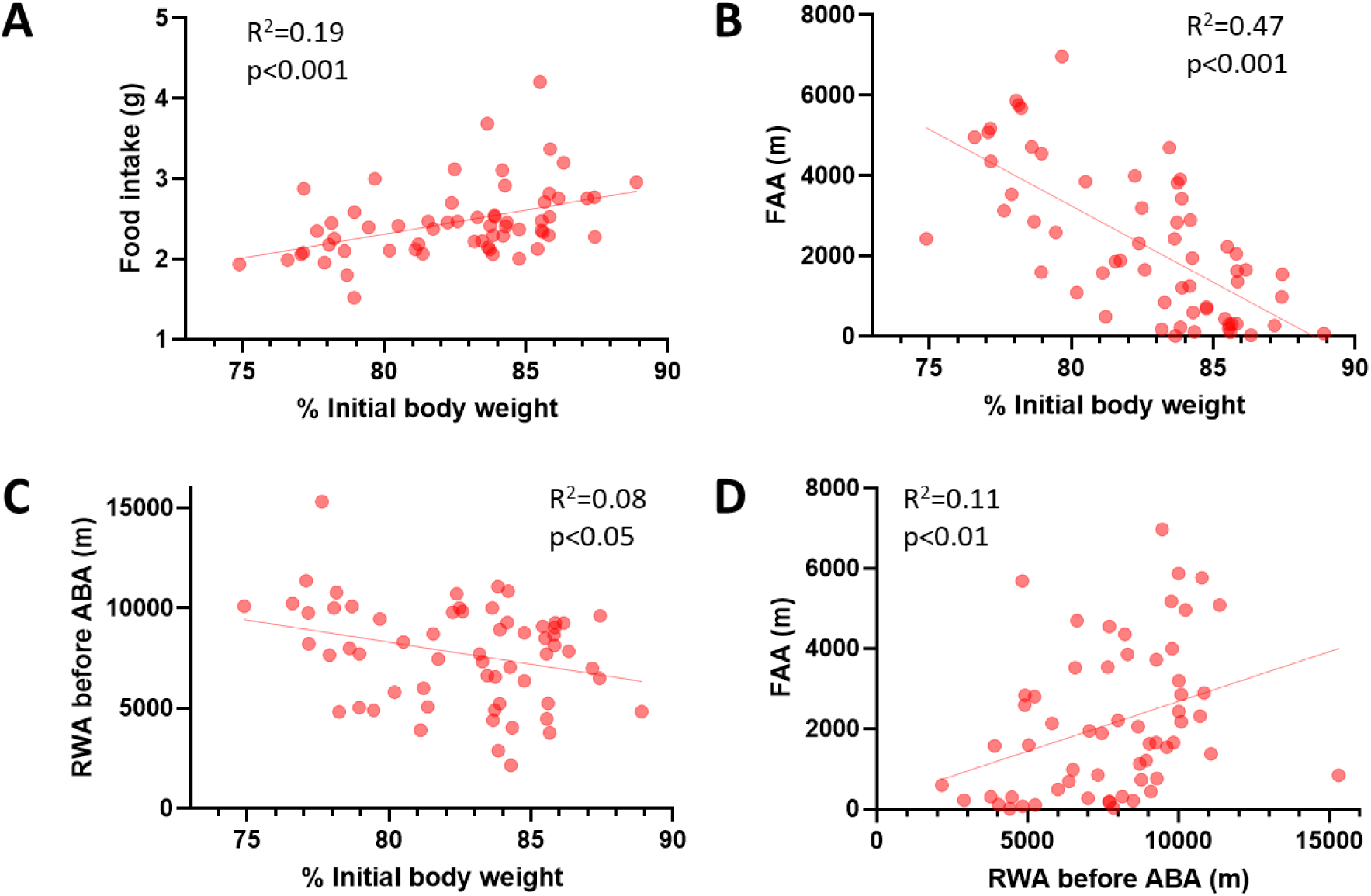
Correlations performed on all ABA-exposed animals used for electrophysiology in this study showing (A) a positive correlation between food intake and body weight loss on ABA day 5, (B) a negative correlation between FAA and body weight loss on ABA day 5, (C) a negative correlation between running wheel activity before ABA exposure and body weight loss on ABA day 5 and (d) a positive correlation between FAA on ABA day 5 and running wheel activity before ABA exposure. *p<0.05, **p<0.01, ***p<0.001. See Table S2 for statistical analyses.

**Supplementary Figure 2: (related to Figure 2).**
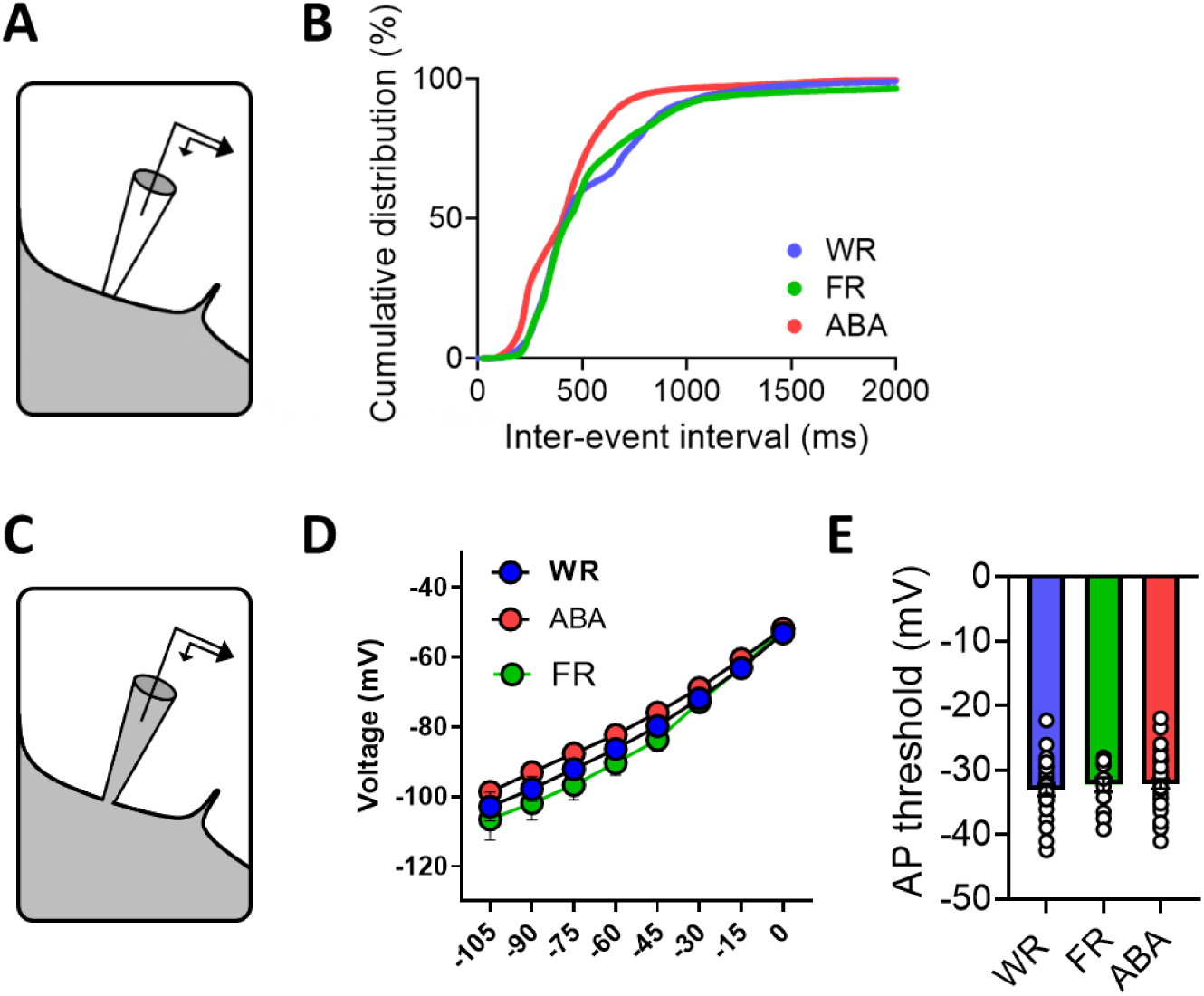
(A) Schematic representation of the cell-attached patch-clamp mode used to record VTADA neurons in Pitx3-GFP mice. (B) Cumulative distribution of action potential inter-event interval. (C) Schematic representation of the whole-cell patch-clamp mode used to record VTADA neurons in Pitx3-GFP mice. (D) Measures of the current-voltage curves. (E) Action potential threshold. Data are means ± SEM. WR, wheel running control group; FR, food restricted control group; ABA, activity-based anorexia group. See Table S2 for statistical analyses.

**Supplementary Figure 3: (related to Figure 3).**
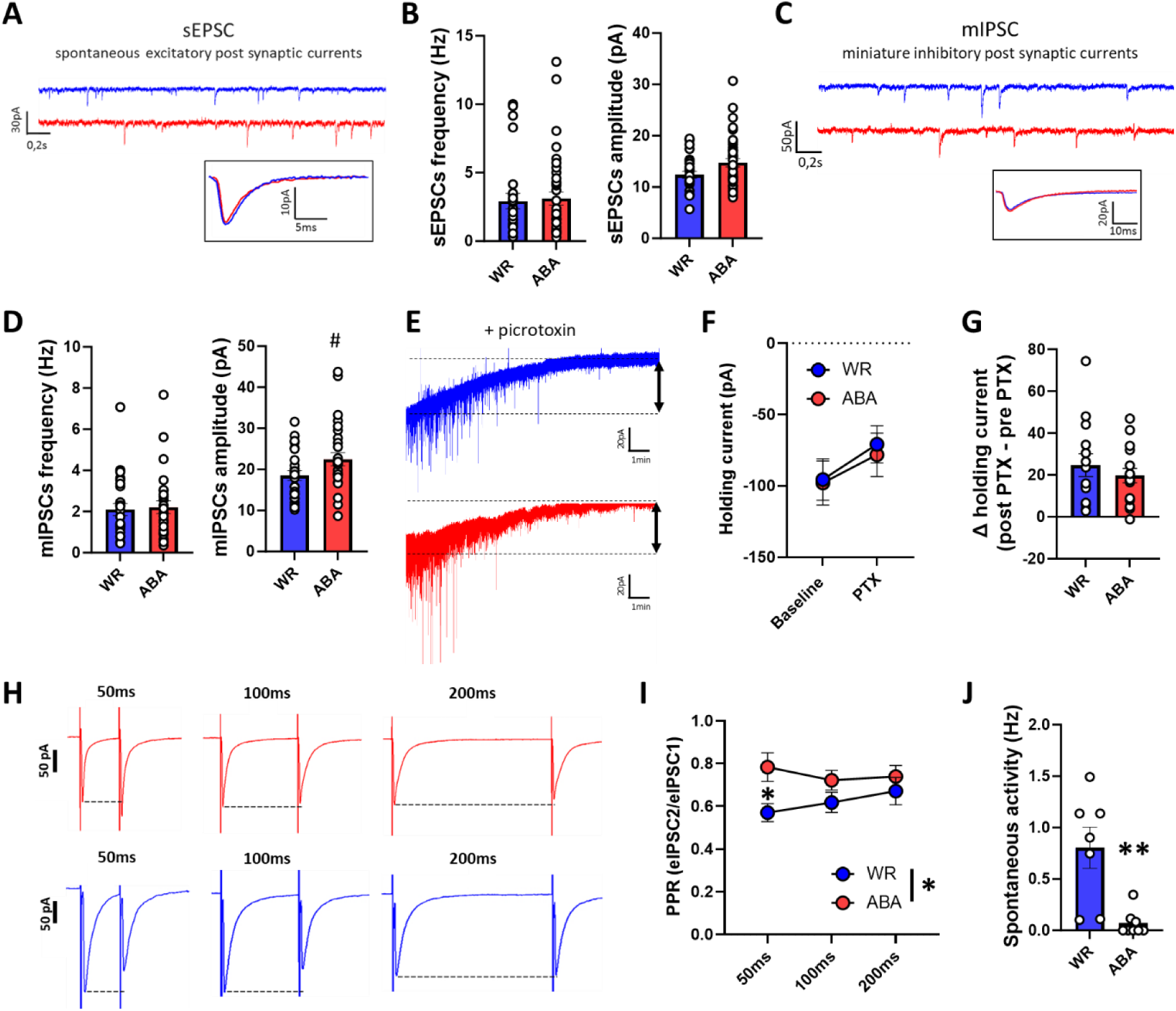
(A) Representative traces of spontaneous EPCSs acquired during voltage clamp recording of VTADA neurons. (B) Measures of sEPSC frequency (left) and amplitude (right). (C) Representative traces of mIPCSs acquired during voltage clamp recording after 1µM TTX application. (D) Measures of mIPSC frequency (left) and amplitude (right). (E) Representative traces and (F-G) collapsed holding current density prior to and following bath application of picrotoxin (100 μM). (H) Example traces of evoked IPSCs with electrical stimulation performed in coronal slices with an inter-stimulation interval of 50ms (left), 100ms (middle) and 200ms (right). (I) Measure of the PPR obtained by dividing the amplitude of eIPSC2 with amplitude of eIPSC1 for each time interval. (J) VTAGABA firing rate averaged for every animal used (the experimental unit is the animal, not the cell). Data are means ± SEM. #p<0.1, *p<0.05, **p<0.01, ***p<0.001. WR, wheel running control group; ABA, activity-based anorexia group. See Table S2 for statistical analyses.

**Supplementary Figure 4: (related to Figure 4).**
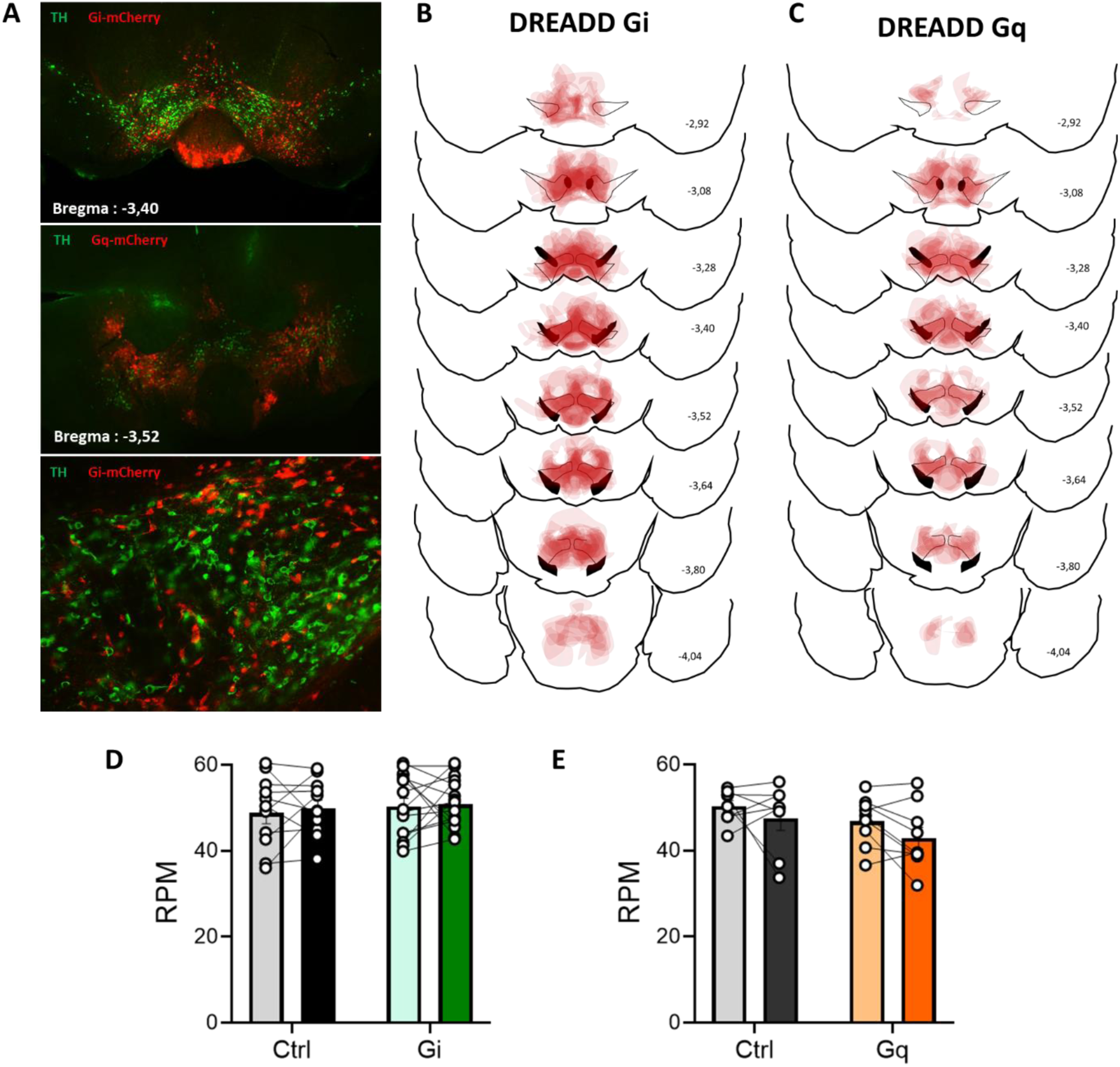
(A) Representative microscopy images showing the expression of DREADD-Gq receptors (top) and DREAD-Gi receptors (middle) mostly in the VTA. The lower image was performed at higher magnification. (B-C) overlay of areas containing DREADD-expressing cells for all animals used in the study for the DREADD-Gi (n=14, C) and the DREADD Gq (n=8, D). (D-E) Rotation per min (RPM) in a rotarod in VGAT-cre mice under inhibition (DREADD-Gi, D) or activation (DREADD-Gq, E) of GABAergic neurons with 0.5mg/kg and 0.25mg/kg of CNO respectively. Data are means ± SEM. TH: tyrosine hydroxylase, Gi: DREADD-Gi receptor-expressing mice, Gq: DREADD-Gq receptor-expressing mice, Ctrl: control group. See Table S2 for statistical analyses.

**Supplementary Figure 5: (related to Figure 5).**
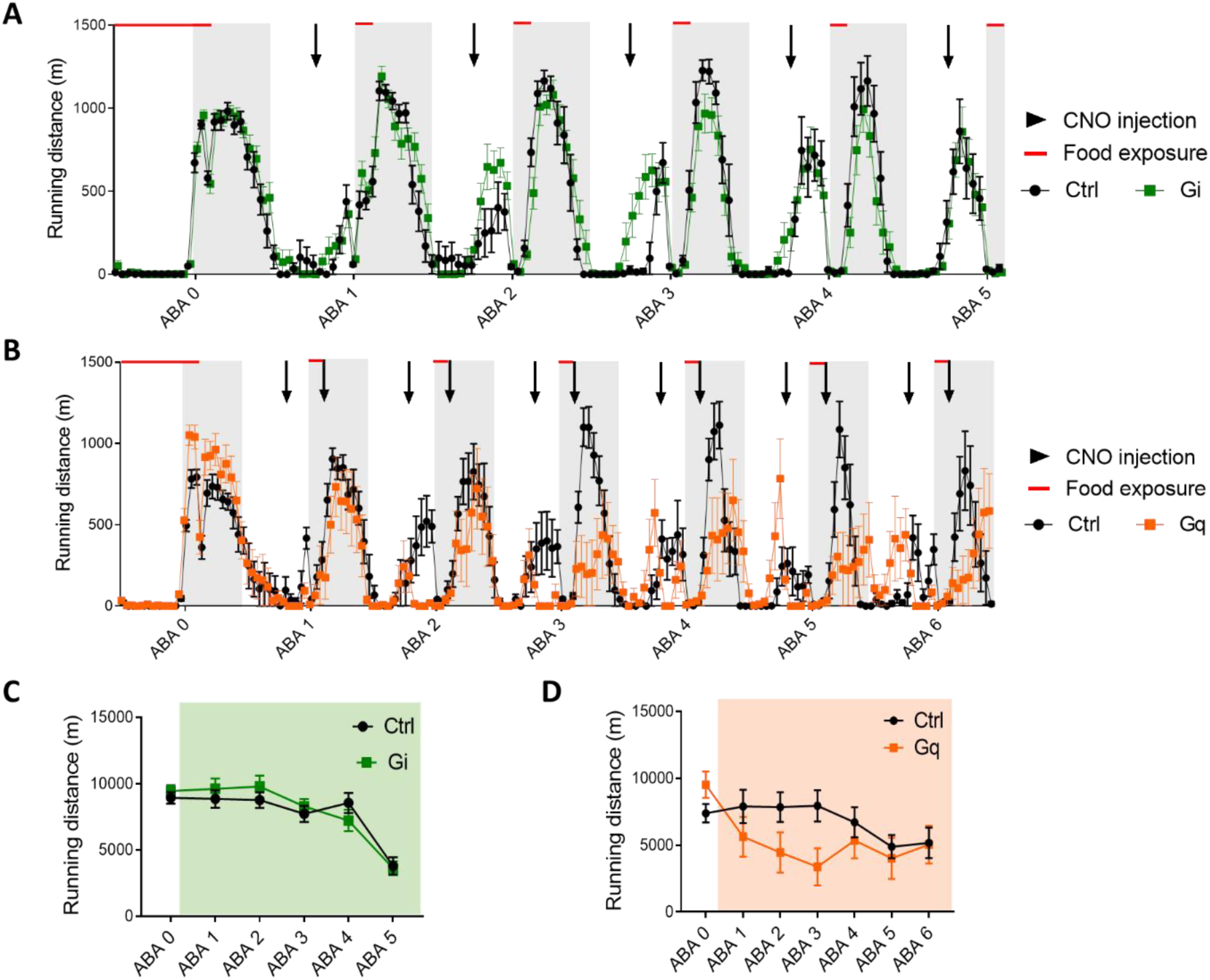
(A) Hour by hour distance travelled in the wheel during ABA day 0 to day 5 with a single 5mg/kg CNO injection per day (black arrows). (B) Hour by hour distance travelled in the wheel during ABA day 0 to day 6 with two 0.25mg/kg CNO injection per day (black arrows). (C) Daily running wheel activity in DREADD-Gi animals for every ABA days. (D) Daily running wheel activity in DREADD-Gq animals for every ABA days. Data are means ± SEM. Gi: DREADD-Gi receptor-expressing mice, Gq: DREADD-Gq receptor-expressing mice, Ctrl: control group, CNO: clozapine-N-Oxide. See Table S2 for statistical analyses.

**Table S2.**
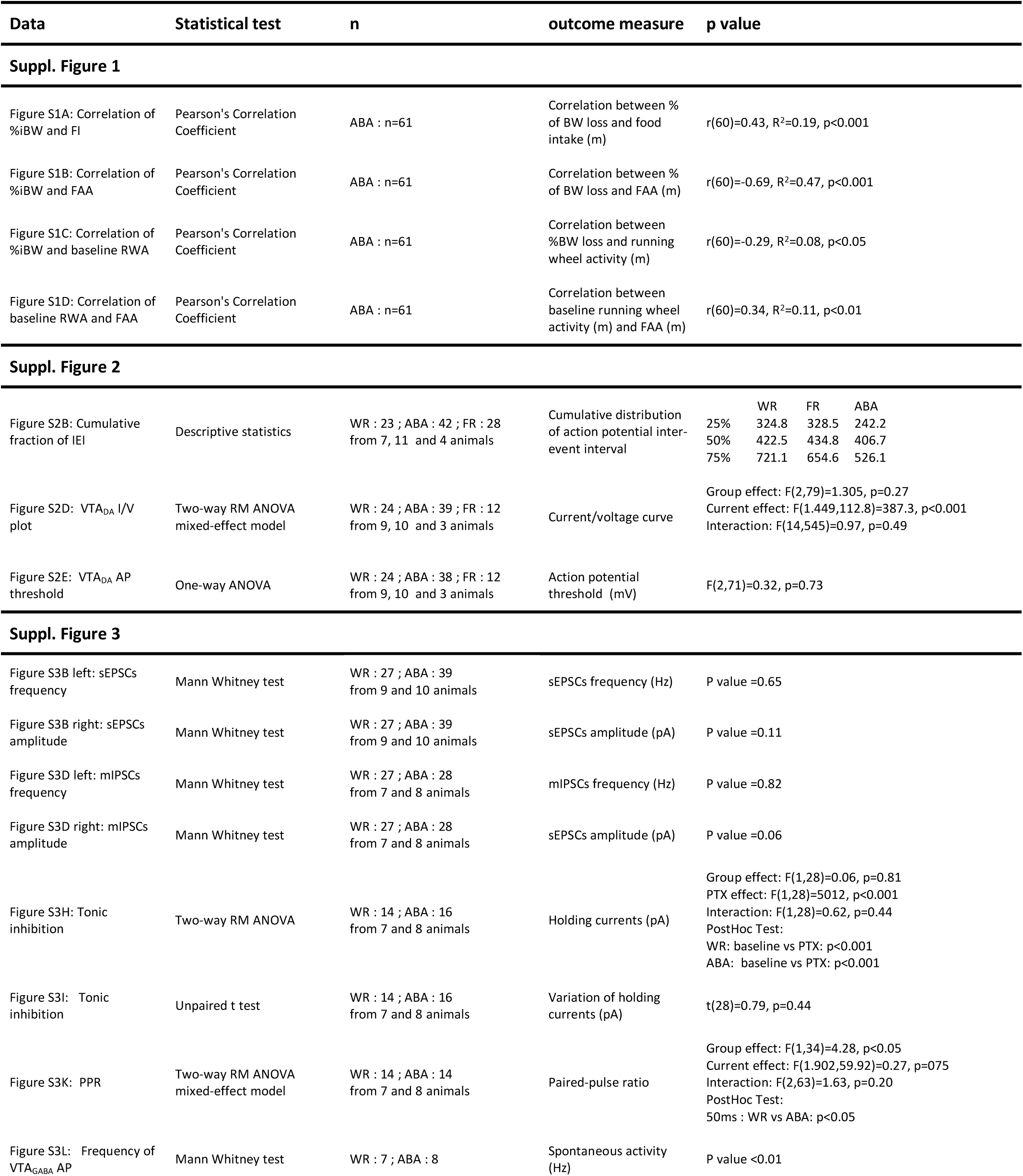
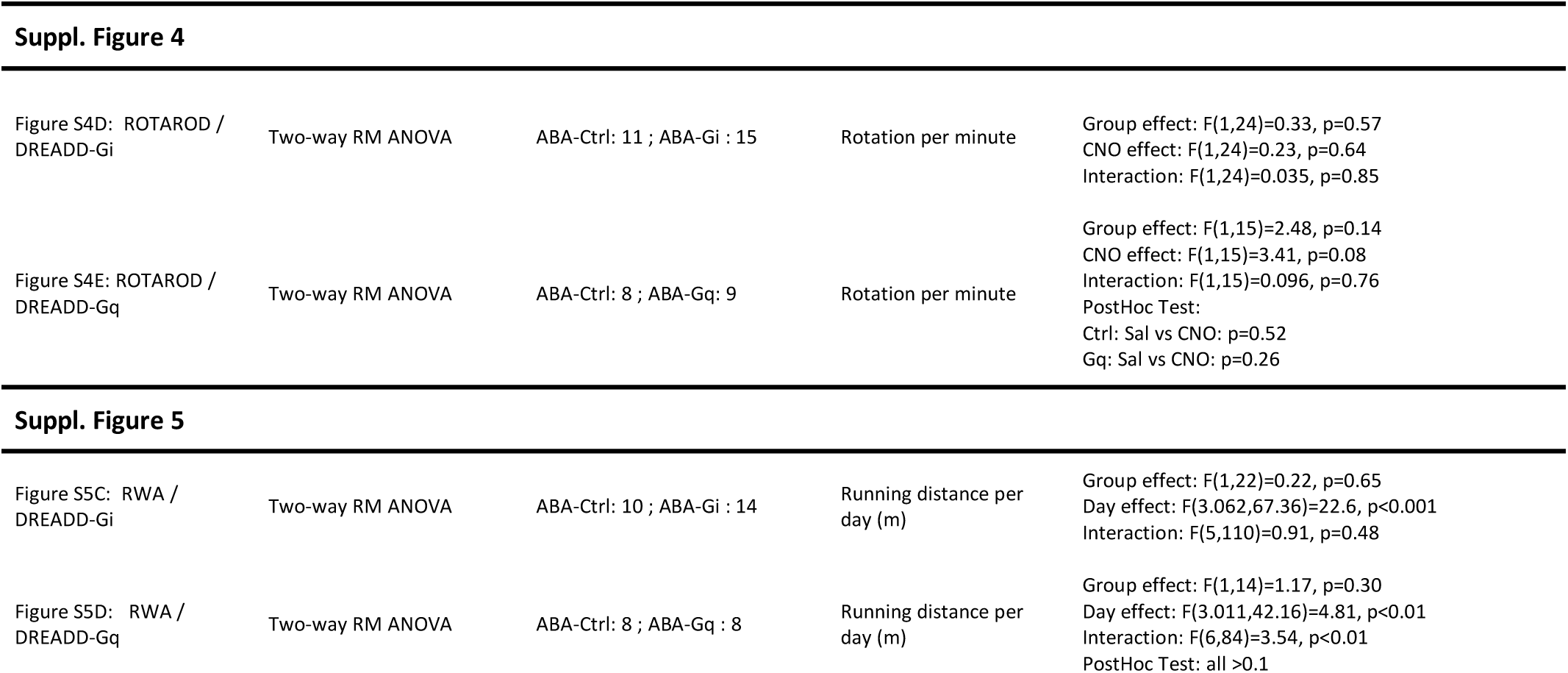
Summary of supplemental statistical analysis

